# Open Science Discovery of Potent Non-Covalent SARS-CoV-2 Main Protease Inhibitors

**DOI:** 10.1101/2020.10.29.339317

**Authors:** Melissa L. Boby, Daren Fearon, Matteo Ferla, Mihajlo Filep, Lizbé Koekemoer, Matthew C. Robinson, The COVID Moonshot Consortium, John D. Chodera, Alpha A Lee, Nir London, Annette von Delft, Frank von Delft

## Abstract

We report the results of the **COVID Moonshot**, a fully open-science, crowd sourced, structure-enabled drug discovery campaign targeting the SARS-CoV-2 main protease. We discovered a non-covalent, non-peptidic inhibitor scaffold with lead-like properties that is differentiated from current main protease inhibitors. Our approach leveraged crowdsourcing, machine learning, exascale molecular simulations, and high-throughput structural biology and chemistry. We generated a detailed map of the structural plasticity of the SARS-CoV-2 main protease, extensive structure-activity relationships for multiple chemotypes, and a wealth of biochemical activity data. All compound designs (>18,000 designs), crystallographic data (>840 ligand-bound X-ray structures), assay data (>10,000 measurements), and synthesized molecules (>2,400 compounds) for this campaign were shared rapidly and openly, creating a rich open and IP-free knowledgebase for future anti-coronavirus drug discovery.

## Introduction

Despite rapid progress in vaccine development, the global failure to abate coronavirus disease 2019 (COVID-19) culminating in more than 690 million confirmed cases worldwide by July 2023, will likely cause the virus to become endemic (*1*), and continue to cause a significant number of deaths, especially in the Global South unless there is an accessible treatment (*2*). Antiviral therapeutics are a necessary and complementary strategy to vaccination in order to control COVID-19(*3*). Several directly acting oral antivirals are now approved for usage against severe acute respiratory syndrome coronavirus 2 (SARS-CoV-2) infection, including ritonavir-boosted nirmatrelvir(*4*), ensitrelvir (Japan)(*5*), and molnupiravir(*6*).

COVID-19 is not an isolated event, but the latest exemplar of a series of threats to human health caused by beta-coronaviruses also responsible for the SARS (2003) and Middle East respiratory syndrome (MERS) (2010) pandemics(*7*). Open knowledge bases and technology infrastructures for antiviral drug discovery will enable pandemic preparedness by refreshing the antivirals pipeline and providing multiple starting points for the development of therapeutics. Here, we report the open science discovery of a potent SARS-CoV-2 antiviral lead compound and a roadmap for the potential development of future SARS-CoV-2 and pan-coronavirus antivirals.

The SARS-CoV-2 main protease (Mpro; or 3CL-protease) is an attractive target for antiviral development due to its essential role in viral replication, a large degree of conservation across coronaviruses, and dissimilarity of its structure and substrate profile to human proteases(*8*) (**FIg S1**). Pioneering studies during and after the 2003 SARS pandemic established the linkage between Mpro inhibition and antiviral activity in cell culture(*9*). This work has been corroborated by in vitro and in vivo studies for SARS-CoV-2(*10*, *11*) and the clinical success of nirmatrelvir (the Mpro inhibitor component of Paxlovid)(*12*) and ensitrevir (Xocova)(*13*, *14*).

To warrant early use in the course of disease or even prophylactically among at-risk populations, an antiviral drug would need to be orally available with an excellent safety profile. Given the historical difficulties in developing peptidomimetic compounds into oral drugs and the risk of downstream idiosyncratic hazards of covalent inhibition, we chose to pursue non-covalent, non-peptidomimetic scaffolds. First-generation oral Mpro inhibitors have now demonstrated clinical efficacy(*15*, *16*), but the need for cytochrome P450 3A4 (CYP3A4) inhibitor co-dosing (ritonavir, in the case of Paxlovid) to achieve sufficient human exposure may substantially limit use in at-risk populations due to drug-drug interactions(*17*). There remains a need for chemically differentiated oral antiviral protease inhibitors with the potential to enter clinical development.

### Crowdsourced progression of X-ray fragment hits rapidly generated potent lead compounds with diverse chemotypes

The COVID Moonshot is an open science drug discovery campaign targeting SARS-CoV-2 Mpro(*18*, *19*), building off a rapid crystallographic fragment screening campaign that assessed 1495 fragment-soaked crystals screened within weeks to identify 78 hits that densely populated the active site (**Figure 1A**)(*20*). This dataset was posted online on 18 Mar 2020(*21*), days after the screen was completed(*21*). The non-covalent fragment hits did not show detectable inhibition in a fluorescence-based enzyme activity assay (assay dynamic range IC_50_ < 100 µM). However, they provided a high-resolution map of key interactions that optimized compounds may exploit to inhibit Mpro(*22*).

**Figure 1:**
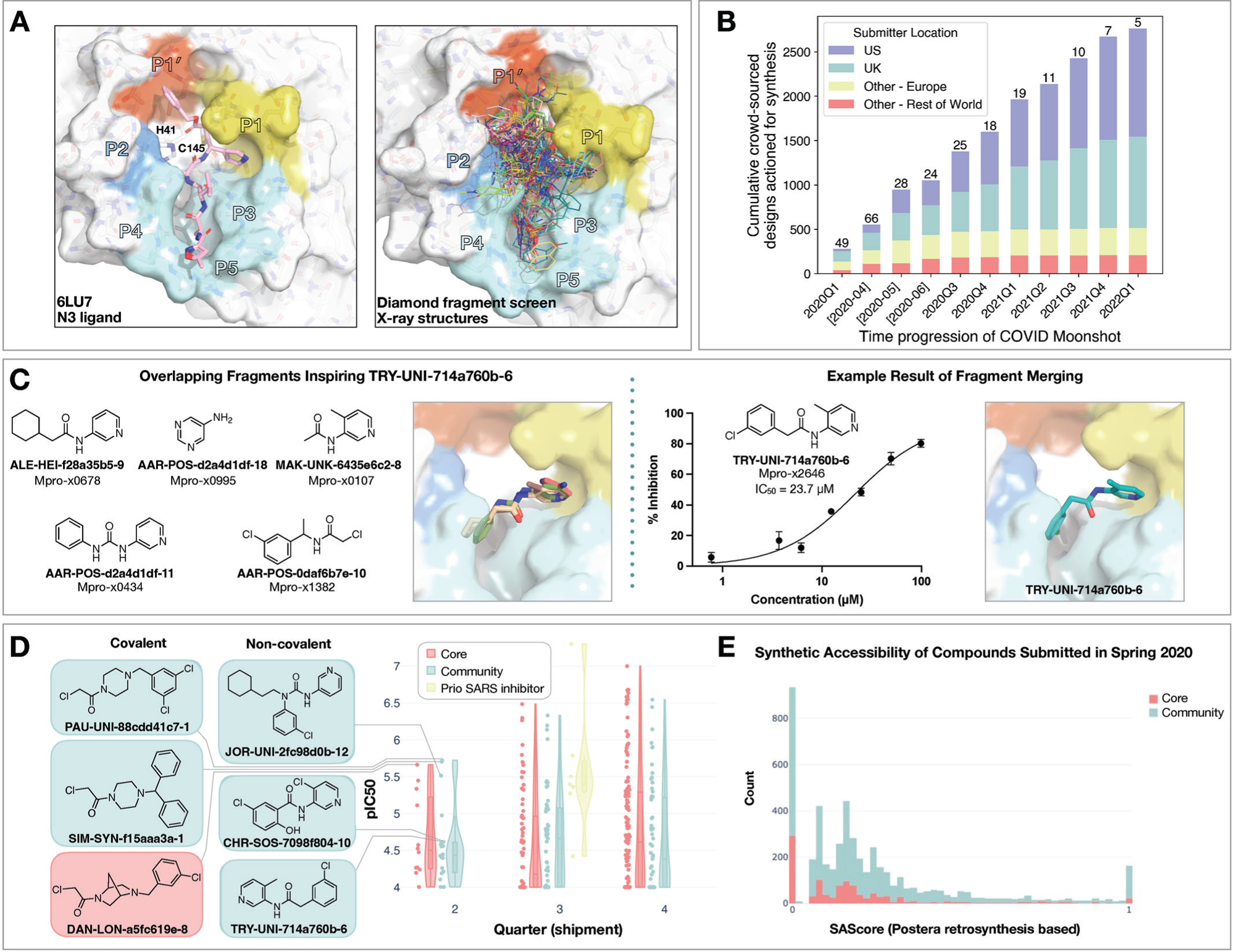
Crowdsourcing rapidly identified novel chemotype scaffolds by merging fragment hits. **A:** A Diamond / XChem fragment screen that initiated this SARS-CoV-2 Mpro inhibitor discovery campaign generated 58 hits that completely cover the Mpro active site, with a variety of chemotypes engaging each pocket; 1495 x-ray datasets were collected and 78 solved structures for hits were publicly posted(*20*). The peptidomimetic N3 ligand is shown at left for comparison to indicate natural substrate engagement in the binding site, defining the peptide sidechain numbering scheme used throughout this work. The nucleophilic Cys145 reacts with the scissile peptide bond between P1 and P1’; His41-Cys145 form a catalytic dyad whose coupled charge states shuttle between zwitterionic and neutral states (*92*). **B:** On March 18th 2020, the COVID Moonshot set up a crowdsourcing website to empower scientists across the globe to contribute molecule designs. The number of designs actioned for synthesis each quarter (except for the 2020Q2 which is shown per-month in brackets) is shown, subdivided by the region of the submitter of the design idea. The total number of unique submitters that contributed actioned designs for that quarter is shown on top of the bars. **C:** Many submissions, such as TRY-UNI-714a760b-6, exploited spatially overlapping fragment hits to design potent leads that are synthetically facile. **D:** Experimental biochemical potency of designs broken down by submission group. Multiple submissions in 2020 from the community were more potent than the best designs from the core team as seen for the top three chloroacetamide structures (left) and non-covalent structures (right). **E:** Distribution of synthetic accessibility scores for designs contributed by the core team and the community. The concern that community submissions may be of poor quality is not supported by the fact that these were as synthetic accessible as those designed by the core team (median: community 0.17 core 0.13,). Half of the outliers (SAScore = 1) were primarily natural products, which are hard to achieve via organic chemistry.

Numerous approaches have been proposed to advance from fragments to lead compounds(*23*, *24*). One strategy, *fragment merging*, aims to combine multiple fragments into a single, more potent molecule, whereas *fragment expansion* elaborates a fragment to engage neighboring interactions. While these strategies are usually applied to a single fragment or a handful of fragments, our large-scale fragment screen produced a dense ensemble of hits, providing an opportunity for rapid lead generation by combining chemotypes from multiple fragments. Nonetheless, this approach requires heuristic chemical reasoning that accounts for the spatial orientation of fragments in the binding site—a feat that can challenge algorithms but is potentially also solvable by humans. Building on successes in crowdsourced protein(*25*) and ribonucleic acid (RNA) (*26*) design campaigns, we hypothesized that crowdsourced human analysis and algorithmic strategies could accelerate the generation of potent lead compounds and furnish diverse chemical matter, as different chemists would employ different approaches and reasoning strategies.

We launched an online crowdsourcing platform [http://postera.ai/covid] on 18 March 2020 (**Figure 1B**), soliciting participants to submit compounds designed based on the fragment hits(*19*). Compounds selected for synthesis were evaluated by biochemical assays (**Data S1**) and x-ray crystallography and the results were released rapidly on the same platform, enabling contributing designers to build on all available data, as well as designs contributed by others. To facilitate transparency and maximal speed, and to avoid delays around intellectual property, all designers were asked to contribute their designs directly into the public domain, with every design and all related experimental data immediately disclosed online, made openly available explicitly free of IP restrictions. This aggressive open science policy enabled contributors from multiple fields in both academia and industry to freely share their ideas. Within the first week, we received over 2,000 submissions, representing a diverse set of design strategies (**Data S2**).

Many submissions exploited spatially overlapping fragment hits. For example, the submission **<u>TRY-UNI-</u> <u>714a760b-6</u>** was inspired by five overlapping fragments, furnishing a non-covalent inhibitor with an SARS-CoV-2 Mpro enzymatic IC_50_ of 23.7 µM (**Figure 1C**). This compound seeded the “aminopyridine” series, whose optimization is described in detail below. We should note that only 11 of the 998 fragments in the DSi-poised library(*27*, *28*) contained a 3-amino pyridine, yet four of them were successfully identified in the structures and were consequently picked-up for merging by the designers. Apart from the aminopyridine series, our campaign identified three other major chemically distinct lead series with measurable potencies against SARS-CoV-2 Mpro inspired by reported SARS-CoV-1 inhibitors (**Fig S2**). Those compounds span the same binding pocket but feature different chemotypes, and the large quantity of structure-activity relationship (SAR) subsequently generated for these series furnishes multiple backup series with different risk profiles. Other groups have subsequently further elaborated on the Ugi(*29*, *30*) and the benzotriazole series we generated(*31*).

Analysis of the submissions provides some hints to the utility of crowdsourcing as a general strategy for hit-discovery or hit-to-lead campaigns. A qualitative assessment of the textual description of submitted designs (**Fig S3**) hints that many of the designers used tools like docking to assess fragment ‘linking’, ‘merging’ or ‘combination’. When trying to more thoroughly categorize submissions it doesn’t appear that hypothesis-driven designs perform better than docking driven designs, however ‘predicting’ historical SARS inhibitors is the best performing strategy (**Fig S4**; **Figure 1D**). Throughout the campaign, designs were contributed by both the core group of labs and medicinal chemists leading this project, as well as by the community. One could hypothesize that the core group being committed to the project, as well as thoroughly invested in the campaign details would contribute more potent designs. However, there is no obvious difference in the distributions of designs produced by the core group vs. the community, in the early stages of the campaign (**Figure 1D**) nor were the designs contributed by the community less synthetically accessible (**Figure 1E**). Later in the campaign, (lead optimization stage) the number of submissions from the community decreased, and comparing potency became irrelevant as other attributes of the molecules were being optimized. It is important to mention that several key compounds along the optimization trajectory (**Figure 5A**) of our lead were contributed by the community and not core group members including: **TRY-UNI-714a760b-6**, **ADA-UCB-6c2cb422-1** and **VLA-UCB-1dbca3b4-15** (the racemic mixture of **MAT-POS-b3e365b9-1**). Although anecdotal, this example demonstrates the potential power of crowdsourcing as a strategy to drive fragment-to-lead campaigns.

### Technologies to support rapid optimization cycles

With a growing number of chemically diverse submissions, we relied on a team of experienced medicinal chemists supported by computational methods to aid in triaging design proposals with the goal of increasing potency. To execute a rapid fragment-to-lead campaign, we used models to plan synthetic routes, enumerate synthetically accessible virtual libraries, and estimate potencies to prioritize which compounds to target for synthesis. We did not use an “autonomous” approach - expert judgment is used to make decisions given all the model predictions. Further, in the context of a fast-moving campaign, we prioritized making progress over granular “human-versus-machine” evaluations.

#### Synthetic route predictions guided decision making to accelerate design-make-test-analyze cycles

We used an established synthetic contract research organization (CRO), Enamine, to carry out rapid synthesis of progressed compound designs. To take full advantage of the available building block collection, we used a machine learning approach that plans efficient retrosynthetic routes to predict synthetic tractability(*32*, *33*). We automatically computed synthetic routes for all crowdsourced submissions utilizing Enamine’s in-stock building block inventories. From the computed routes, synthetic complexity was estimated based on the number of steps and the probability of success of each step. The synthetic accessibility score, as well as the predicted synthetic route, were then used to aid medicinal chemistry decision making. Our predicted synthetic complexity correlated with the actual time taken to synthesize target compounds and the algorithm was able to pick out advanced intermediates as starting materials (**Figure 2A**).

**Figure 2:**
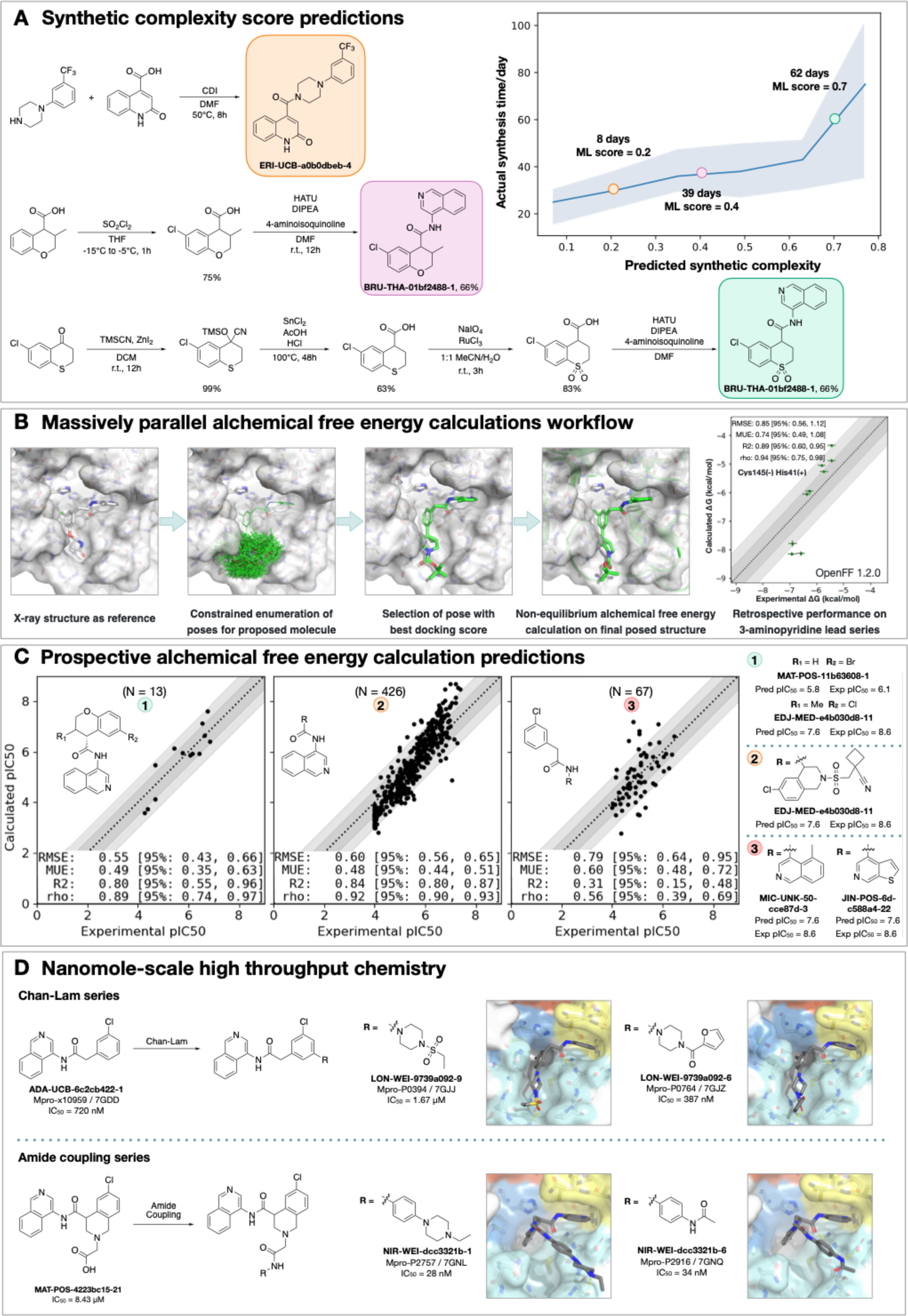
Strategies to support rapid optimization cycles. **A:** Machine learning forecasts experimental synthesis time (left) and returns efficient routes that leverage over 10 million in-stock advanced intermediates (right). Our algorithm predicts the probability of each step being successful, and predicts synthetic accessibility by taking the product of the probabilities along the whole route. We analysed all compounds made in COVID Moonshot from 2020-05-01 to 2021-07-01 (n=898). The right panel exemplifies the experimental execution of the predicted routes, demonstrating the ability of the algorithm to build on functionalized intermediates to shorten synthesis. **B:** Applying alchemical free energy calculations at scale enables us to estimate the potency of compounds. Retrospective assessment of our automated free energy calculation workflow on early compounds in the 3-aminopyridine series in the first month of the COVID Moonshot campaign suggested free energy calculations could provide good predictive utility, which inspired confidence for large scale deployment during this campaign. Here, the absolute free energy of binding ΔG is shown in the rightmost panel by adding a constant offset to the computed relative free energy differences. **C:** Alchemical free energy predictions for all submissions elaborating on the depicted scaffold for three representative batches of prospective free energy calculations plotted as calculated (converted using Cheng-Prusoff equation) versus experimental pIC_50_. Simulations were run using Mpro in dimer form, with neutral catalytic dyad and no restraints. Each batch (numbered 1-3 from left to right) is annotated with its scaffold and top-scoring candidates are shown on the right-hand side (numbered 1-3 from top to bottom)---for these the structure names are shown together with their predicted and experimental pIC_50_ (“Pred”/”P” and “Expt”/”E”, resp.). Statistical performance with 95% confidence intervals for each batch is shown as a table in each scatterplot. **D:** Two examples of nanomole scale high throughput chemistry (HTC) campaigns used to optimize the potency of intermediate binders, centering on the Chan Lam reaction (**Fig S7**) and amide couplings (**Fig S8**). Direct biochemical screening of crude reactions identified candidates that were resynthesized and in both cases were able to improve the potency of the parent compound. Soaking of crude reaction mixtures of the most potent biochemical hits into Mpro crystals provided complex structures with the identified hits (Chan-Lam - PDBs: 7GJJ/7GJZ resolution: 1.75Å/1.65Å; Amide coupling PDBs: 7GNL/7GNQ resolution: 1.68Å/1.53Å). In both cases new interactions were discovered, explaining the improved activity. It is interesting to note, that while for the Chan Lam reaction campaign, the extended compounds indeed occupied the intended P4, for the amide-coupling vector, all compounds extended into the P3/5 pockets.

#### Alchemical free energy calculations prioritized potent compounds for synthesis

We estimated potency of proposed designs and virtual synthetic libraries of analogues using alchemical free energy calculations(*34–36*), an accurate physical modeling technique that has hitherto not been deployed in a high throughput setup due to its prohibitive computational cost. We employed Folding@Home(*37*)—a worldwide distributed computing network where hundreds of thousands of volunteers around the world contributed computing power to create the world’s first exascale computing resource(*38*)—to compute the free energy of binding of all 20,000+ crowdsourced and internal design submissions using the Open Force Field Initiative “Parsley” small molecule force fields(*39*) and nonequilibrium switching with the open source PERSES alchemical free energy toolkit(*40–42*) based on the GPU-accelerated OpenMM framework(*43*)(*38*) (see **Detailed Methods**). Comprehensive sampling was prioritized over efficiency of computation given the abundant compute resources available on Folding@home.

We first performed a small retrospective study using bioactivity data generated from the first week of crowdsourced compound designs, triaged solely using synthetic accessibility. The results of these free energy calculations showed good correlation with experimentally-measured affinities (**Figure 2B**). Henceforth, alchemical free energy calculations were used as an additional (though not sole) criteria to guide compound selection and iterative design (see **Data Availability**). During the campaign, distinct objectives were solicited from submitters to address medicinal chemistry problems, and free energy calculations were used to assess these submissions based on predicted potency. **Figure 2C** shows that predicted pIC_50_ tracks experimental measurements across 3 chronologically distinct design campaigns: decoration of the benzopyran ring, replacement of the benzopyran system, and replacement of the isoquinoline system. Note that some design ideas with low predicted pIC_50_ were synthesized since the medicinal chemistry team balanced between gaining insights on structure-activity and structure-property relationship and potency optimization. The champion compounds from each design campaign are highlighted on the right panel of **Figure 2C**. Although free energy calculations identified multiple potency-improving transformations, the strategically useful one was the swap from pyran to a piperidine sulfonamide system, which is on the critical path to the lead compound (**Figure 5A**). On average, 80 GPU hours/compound was used across the three panels (see **Detailed Methods**).

A major strength of alchemical free energy calculations proved to be their ability to select potent analogues from virtual synthetic libraries from which the medicinal chemistry team had already selected compounds sharing a common intermediate, as well as highlighting submitted designs predicted to be highly potent but where major synthetic effort would be required. Our design team prioritized for synthesis small libraries suggested by the aforementioned computational approaches. Chemically-related groups of outliers frequently provided new chemical insight that informed modeling choices (**Fig S5**). The approach was not without drawbacks, including the need to focus on a single reference compound and structure to design transformation networks (rather than leveraging the abundant structural data), the requirement that protonation states be fixed for the entire calculation (requiring the entire transformation network to be recomputed to assess a different protonation state), and the relatively large computational cost required to handle large replacements (see **Detailed Methods**). The method is also not uniformly accurate across all chemical transformations, and accurately estimating its accuracy beforehand is challenging. For example, isoquinoline replacements show lower correlation between calculated and predicted free energy (Panel 3 of **Figure 2B**) compared to the benzopyran replacements (Panel 2 of **Figure 2B**).

#### Nanomole-scale high-throughput chemistry enabled rapid evaluation of structure-activity relationship (SAR)

A complementary method for rapid SAR evaluation was the use of nanomole scale high-throughput chemistry(*44*, *45*) (HTC), coupled with a ‘direct to biology’(*46–48*) biochemical screening. Two examples include the optimization of the Chan-Lam reaction(*49*) to extend molecule **ADA-UCB-6c2cb422-1** and amide coupling to extend **MAT-POS-4223bc15-21** (**Figure 2D**). In both cases, we determined the co-crystal structures of the parent compounds (**Fig S6**) and suggested vectors that could target the P4 pocket of Mpro. Optimization of the reaction conditions was performed for the starting building block with model amines (**Figs S7-S8**) and the optimal conditions were applied to HTC with a library of 300 amine building blocks (**Data S3**). Yield estimation was performed in both cases and showed for the Chan-Lam library only 29 of the library yielded >30% of the desired product, in comparison to 151 for the amide coupling. Nevertheless, the crude mixtures were subjected to a biochemical assay against Mpro (**Data S3**). Seven compounds were selected for resynthesis from the Chan-Lam series and 20 from the amide series (**Fig S9**). In parallel to synthesis, the crude reaction mixtures were subjected to soaking and x-ray crystallography. The structures verified the extended compounds indeed adopt a similar binding mode to the parent. Chan-Lam extended compounds indeed occupied P4, whereas the amides extended towards P3/P5, in both cases forming new interactions with Mpro (**Figure 2D**). Upon resynthesis, one of the Chan-Lam compounds was able to slightly improve over the parent compound IC_50_. Several of the amide-coupling series were able to improve by up to 300-fold on the parent acid-compound (up-to 3-fold on the corresponding methyl-amide) with the best inhibitor exhibiting an IC_50_ of 28nM against Mpro.

#### Covalent targeting strategies

Another approach that was attempted in order to rapidly gain potency was the use of electrophiles to covalently target the catalytic Cys145. The original fragment screen(*20*) that launched this effort, included electrophiles(*50*) and resulted in 48 structures of covalently bound fragments, the vast majority of which were chloroacetamides. Some of the earliest, and most potent, fragment merges explored by both the core group as well as the community were of chloroacetamide (**Figure 1D**) and further optimization improved chloroacetamide fragments’ IC_50_ to as low as 300nM (**Fig S10**). Chloroacetamides, however, are not considered suitable for therapeutics and therefore we aimed to move towards acrylamides by derivatizing potent reversible hits(*30*) (**Fig S11**). Ultimately, we focused on a non-covalent series, but the chlorophenyl moiety that remained throughout the series (**Figure 5A**) was adopted from a chloroacetamide hit fragment (**AAR-POS-0daf6b7e-10; Figure 1C**).

### High-throughput structural biology uncovered binding modes and interactions underlying potency

We selected compounds based on synthetic tractability and alchemical free energy calculations. We profiled every compound through crystal soaking and x-ray diffraction totaling in 584 structures (see **Table S1 and Fig S12** for average statistics; **Data S4** for Crystallographic and refinement statistics and **Fig S13** for ligand density for the structures highlighted in this manuscript). Analysis of a subset of this large trove of structural data, (n=367, up to July 2021), reveals the hotspots for ligand engagement and plasticity of each binding pocket. **Figure 3** highlights the statistics of intermolecular interactions between the residues and our ligands. The P1 and P2 pockets are the hotspots of interactions, yet the interaction patterns are starkly different. The salient interactions sampled by our ligands in the P1 pocket are H163 (H-bond donor), E166 (H-bond acceptor), and N142 (hydrophobic interactions). Whereas P2 interactions are dominated by π-stacking interactions with H41 and hydrophobic interactions M165. The P1’ and P3/4/5 pockets are sparingly sampled by our ligands; the former can be targeted via hydrophobic interactions (T25), whereas the latter via H-bonds (Q192).

**Figure 3:**
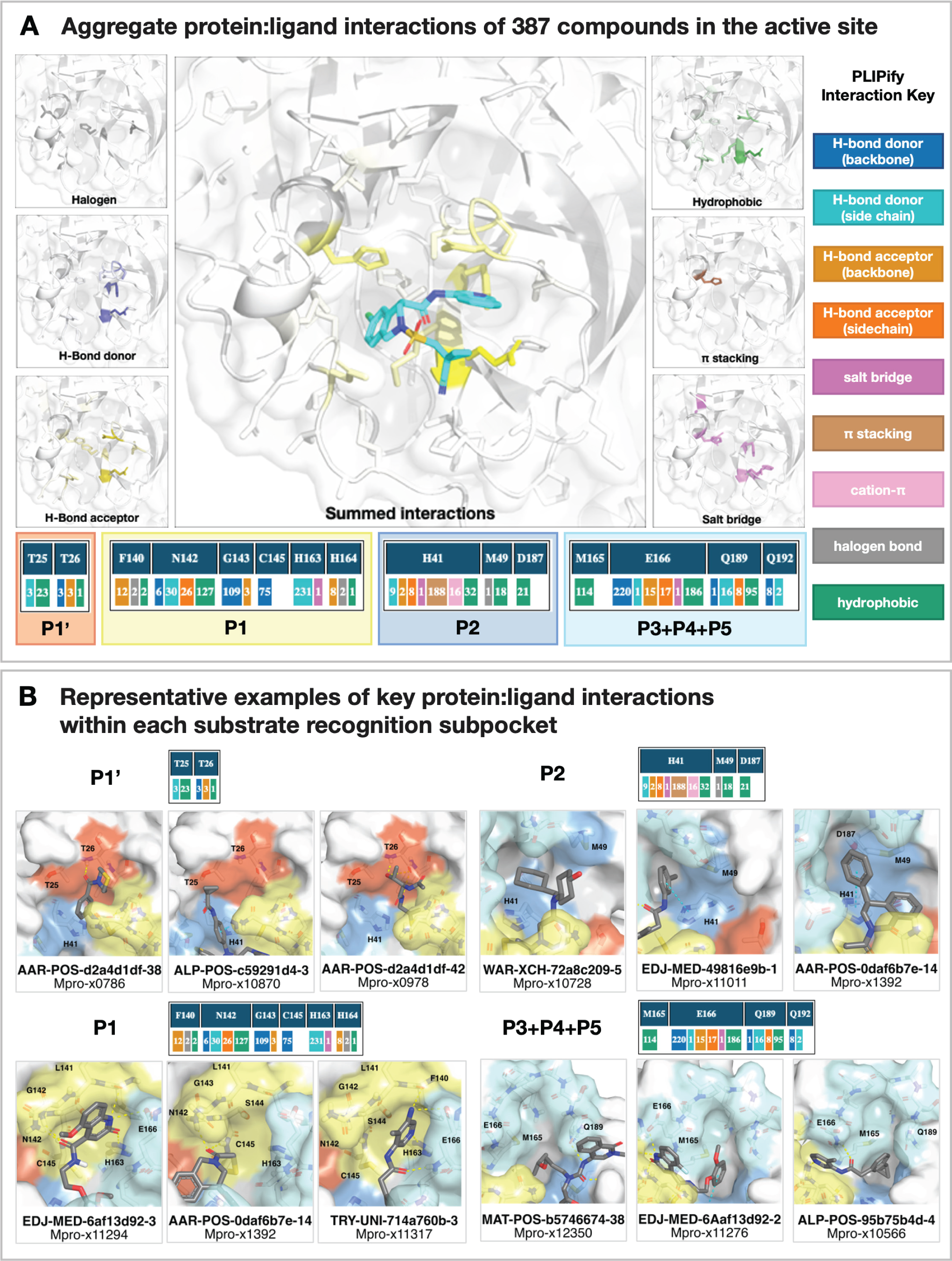
Analysis of 367 complex crystal structures reveals hotspots for ligand engagement, and a variety of ways to engage substrate recognition subpockets. **A:** The five substrate recognition subpockets exhibit distinct preferences for intermolecular interactions. The figure highlights the locations of different types of interactions, with the shading indicating the frequency. The bottom row tallies the number of times each interaction was seen in our structures for different residues. The interaction map was generated using PLIPify (see Detailed Methods) and summarizes the interactions witnessed across 367 complexes from the perspective of the protein, distinguishing between backbone (bb) and sidechain (sc) interactions (which might be more vulnerable to point mutations). **B:** Representative examples of protein-ligand interactions engaging the P1’, P1, P2, and P3-5 subpockets. Hydrogen bonds and π-stacking interactions are depicted as yellow and cyan dashed lines, respectively. The rows above each set of complexes tally the number of times each interaction was seen with the specific residues within the subpockets. See **Data S4** for PDB IDs and crystallography statistics.

This pattern of intermolecular interactions is reflected in the plasticity of the different subpockets. The dominance of directional interactions in P1 renders it more rigid than P2 (**Figure 4**). The rigidity is also dependent on the chemical series (**Fig S2**), with the Ugi and benzotriazole series being able to deform the P2 pocket. Those series comprise more heavy atoms and span a larger region of the binding site, thus changes in P2 pocket interactions could be better tolerated.

**Figure 4:**
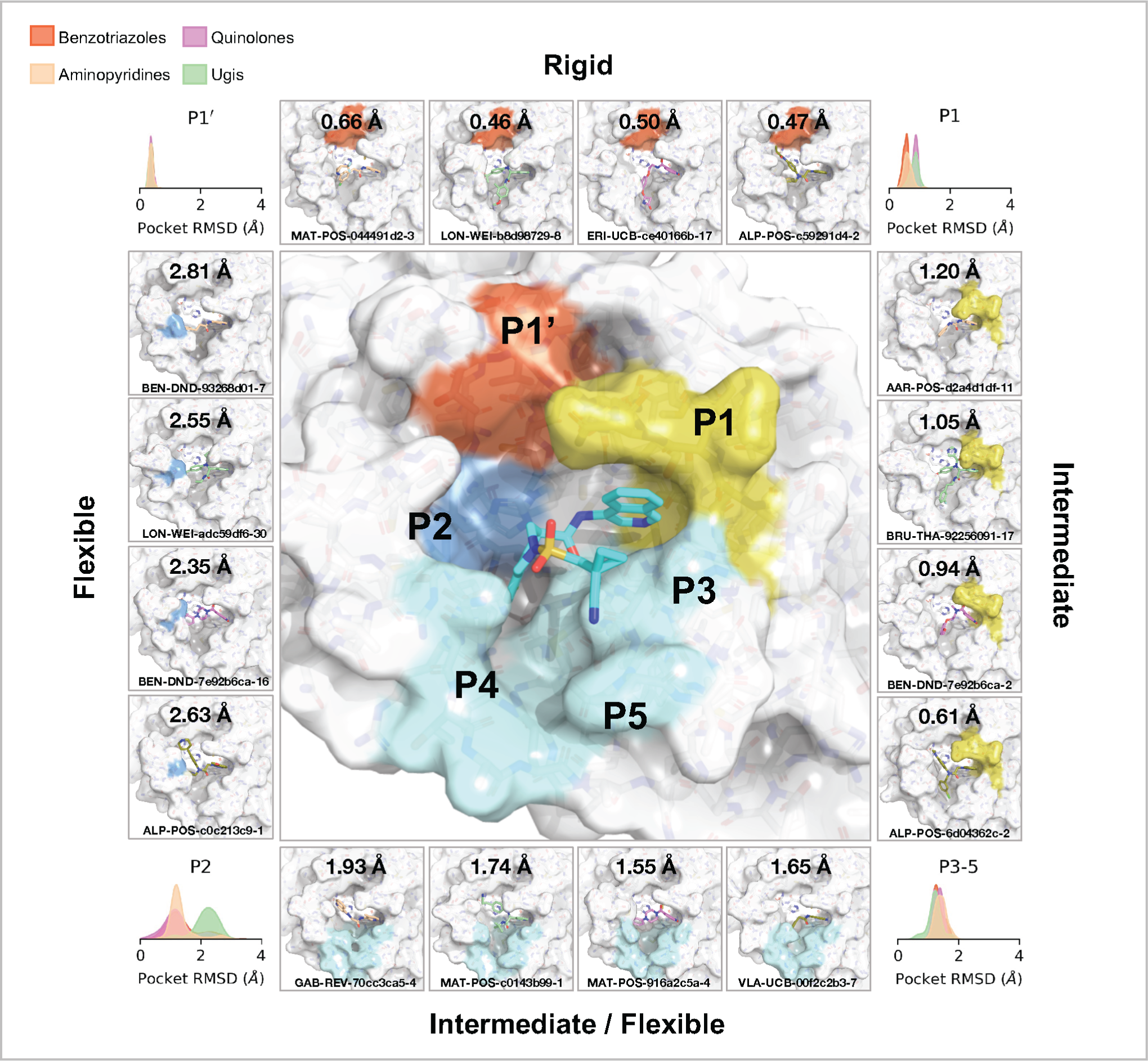
Structural plasticity of the binding subpockets. The subpockets have different degrees of plasticity, which is also dependent on the chemical series (**Fig S2**). The corners of the figure shows the distribution of side chain root-mean-square-deviation (RMSD) deviations from the structure of **MAT-POS-e194df51-1** (middle panel; PDB: 7GAW). The boxes exemplify ligands that significantly deform the pockets.

### Design of a SARS-CoV-2 Mpro inhibitor lead series with potent antiviral activity

Our medicinal chemistry strategy was driven by the design of potent ligand-efficient and geometrically compact inhibitors that fit tightly in the substrate binding pocket. The former strategy aimed to increase the probability of achieving oral bioavailability, while the latter heuristic was motivated by the substrate envelope hypothesis for avoiding viral resistance(*51*). **Figure 5A** outlines the critical intermediates on the path towards an optimized lead compound.

**Figure 5:**
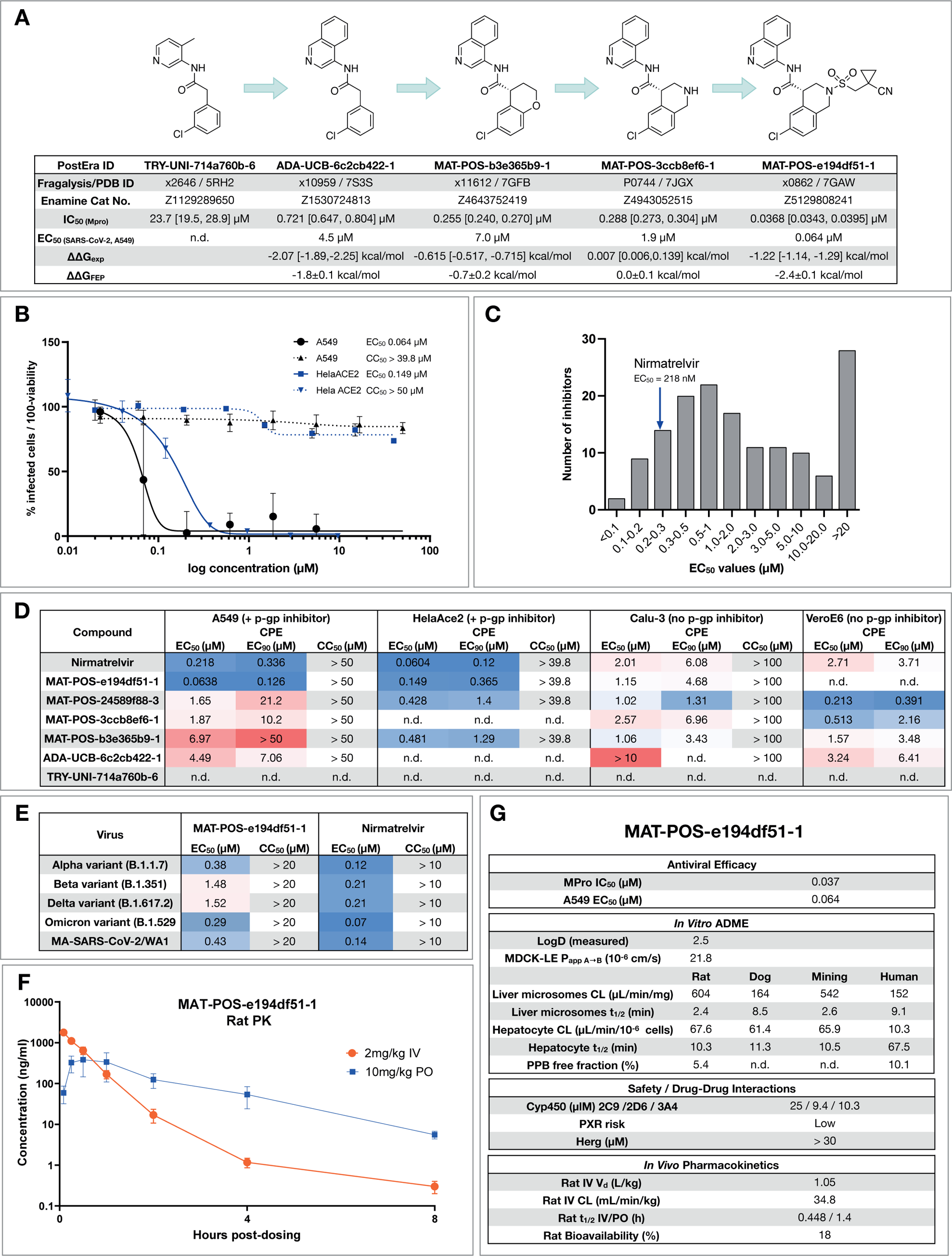
Iterative medicinal chemistry furnished an orally bioavailable inhibitor. **A:** Summary of key medicinal chemistry milestones in developing the initial crowdsourced lead compound into a potent antiviral. X-ray structures for each milestone compound are available via Fragalysis, and each compound can be obtained from Enamine via the corresponding catalogue numbers. Retrospective alchemical free energy calculation predictions for each transformation (ΔΔG_FEP_) are shown for each step between milestones, along with the corresponding experimental free energy difference (ΔΔG_exp_) derived from biochemical activities. As positive control, under our assay condition Nirmatrelvir has IC_50_ of 2.6nM. **B:** Antiviral activity of **MAT-POS-e194df51-1** cellular antiviral assays, with an EC_50_ of 64 nM in A549-ACE2-TMPRSS2 cells assessing cytopathic effect (CPE, black, plotted as 100 - % viability), and 126 nM in HelaAce2 assays (blue, plotted as % infected cells). Both assays were performed with p-gp inhibitors. **C**: Histogram comparing antiviral efficacy of all COVID Moonshot compounds measured to date in an A549-ACE2-TMPRSS2 CPE cellular antiviral assay. **D:** Detailed cellular antiviral assessment of key compounds comprising the synthetic strategy (Fig 5A) across different cell lines and assay techniques, with and without p-gp inhibitors, demonstrating efficacy of **MAT-POS-e194df51-1** in various set-ups and laboratories. **E**: **MAT-POS-e194df51-1** shows good cross-reactivity against known circulating variants of SARS-CoV-2 in antiviral cellular assays in a CPE assay in HelaACE2 cells. **F**: PK profile of **MAT-POS-e194df51-1** in rats with a 2 mg/kg intravenous and 10 mg/kg oral dosing with good oral availability. **G**: ADME characteristics of **MAT-POS-e194df51-1** demonstrate a favorable safety profile, indicating translational potential of the lead series.

Starting from the fragment hit, we explored the P1 pocket, which admits a steep SAR, perhaps unsurprising given its rigidity and preference for directional H-bond interactions (**Figure 3A**). An increase in potency was unlocked by replacing pyridine with isoquinoline, which picks up additional hydrophobic interactions with N142. The SAR around the P2 pocket is considerably more tolerant to modifications, and broadly favors hydrophobic moieties. A step-change in potency was achieved by rigidifying the scaffold: We introduced a tetrahydropyran ring to transform the P2 substituent into a chromane moiety (compound **MAT-POS-b3e365b9-1**; the racemic mixture **VLA-UCB-1dbca3b4-15**, which was initially synthesized, has a IC_50_ of 360 nM; **Figure 5A**), chosen because of building block availability. Despite possessing a degree of molecular complexity, **MAT-POS-b3e365b9-1** is obtained via a one-step amide coupling (**Figure 2A**). We then further explored the P2 pocket with a library chemistry strategy in mind. Thus, guided by free energy calculations (**Figure 2C**), we first substituted the chromane for a tetrahydroisoquinoline to introduce a functionalizable handle (**MAT-POS-3ccb8ef6-1, Figure 5A**), which maintained potency. Finally, we constructed a focused library realized via sulphonamide Schotten-Baumann coupling (**Fig. S14**), furnishing an increase in both enzymatic inhibition and cellular antiviral efficacy. This work led to a potent antiviral chemical series (**Figure 5A**) with a favorable safety profile, low brain penetrance (**Fig S15; Data S5**), improved oral bioavailability, but moderate in vitro-in vivo correlation in clearance (**Fig S16; Data S5**; all measured cellular antiviral data is available in **Data S6**).

As an example for the aminopyridine lead series, we discuss antiviral efficacy, absorption, distribution, metabolism and excretion (ADME) and pharmacokinetic (PK) characteristics of compound **MAT-POS-e194df51-1:**. **MAT-POS-e194df51-1** was profiled in SARS-CoV-2 antiviral assays across multiple cell lines,exhibiting EC_50_ of 64 nM in A549-ACE2-TMPRSS2 cells and 126 nM in HelaAce2, without measurable cytotoxicity (**Figure 5B**). This is in line with overall cellular efficacy for the chemical series; of 150 compounds with enzyme assay IC_50_ <500 nM assessed in A549-ACE2-TMPRSS2 cellular cythopatic effect (CPE) assays, 15 compounds showed lower EC_50_ values compared to the internal control nirmatrelvir that was measured at an EC_50_ of 218 nM in this assay (**Figure 5C**). Similarly, good antiviral activity was measured across “crowdsourced” antiviral assays across different laboratories and cell lines, including assays performed with and without p-gp inhibitors and using nirmatrelvir as an internal control (**Figure 5D**). We also observed good cross-reactivity of our lead compound **MAT-POS-e194df51-1** against known SARS-CoV-2 variants Alpha, Beta, Delta and Omicron (**Figure 5E**). Closely related molecules **PET-UNK-29afea89-2** and **MAT-POS-932d1078-3** with EC_50_s in HelaAce2 CPE assays of 240 nM and 331 nM and 657 nM and 2.57 µM in A549-ACE2-TMPRSS2 CPE assays, respectively (**Fig S17A** and **B**) show a >100-fold reduction of intracellular viral RNA and infectious virus secretion into the apical compartment of human induced pluripotent stem cell-derived kidney organoids (**Fig S16D** and **E**), an accessible model for the human kidney, an organ that is infected in COVID-19 patients, as reported previously for earlier analogues of the same series(*52*). **MAT-POS-e194df51-1** exhibits favorable properties required for an orally bioavailable inhibitor (**Figure 5F,G**). In addition, crystallographic studies reveal that the interaction pattern of **MAT-POS-e194df51-1** with the Mpro binding site is distinct to approved Mpro inhibitors nirmatrelvir and ensitrelvir (S-217622) (**Fig S18)**, thus potentially offering complementary resistance profiles and justifying further development.

### Open Science presents a viable route to early drug discovery

The results presented here reflect the success of an open science, patent-free antiviral discovery program in rapidly developing a differentiated optimized lead in response to an emerging pandemic threat. As a result of the open science policy, a large number of collaborators (now **The COVID Moonshot Consortium**) were able to provide in-kind support, providing synthesis, assays and in vitro/vivo experiments. By making all data immediately available, and all compounds purchasable from Enamine, we aim to accelerate research globally along parallel tracks following up on our initial work. As a striking example for the impact of open-science, the Shionogi clinical candidate S-217622 (which has now received emergency approval in Japan as Xocova [ensitrelvir]) was identified in-part based on crystallographic data from The COVID Moonshot Consortium(*53*).

Despite our optimization and characterization efforts, considerable gaps from reporting a clinical candidate remain: The series requires further PK and pharmacodynamic (PD) optimization; in particular it displays high clearance and low bioavailability. As it stands, it would likely not be able to achieve therapeutic exposure without a pharmacokinetic booster (such as ritonavir). To move forward, additional in-depth safety data is required, as well as additional PK data from a second species to enable accurate human dose prediction. The COVID Moonshot and its lead series for COVID-19 have been adopted into the drug development portfolio of the Drugs for Neglected Diseases Initiative (DNDi) for further lead optimization and downstream pre-clinical development. This work is funded by a $10M Wellcome Trust through the COVID-19 Therapeutics Accelerator (ACT-A), of which results will be reported upon filing Clinical Trials Authorization (CTA) (*54*). To reach Phase II readiness we expect a further $7.5M will be required in addition to process route development costs(*55*).

Open science efforts have transformed many areas of biosciences, with examples such as the Human Genome Project(*56*), the Structural Genomics Consortium(*57*), and the RAS Initiative(*58*). The COVID Moonshot provides an example of open science drug discovery leading to advances in infectious diseases drug discovery—a research area of grave public importance but one which is chronically underfunded by the private sector(*59*).

## Methods Summary

### 0. Compound registration and data flow process

All compound designs from the internal medicinal chemistry team, collaborators, and external submitters were captured through the online compound design submission platform [https://postera.ai/covid] along with submitter identity, institution, design rationale, and any inspiration fragments. A forum thread was created to discuss these designs and attached to the compound design. Each submitted batch of related designs received a unique ID including the first three letters of the submitter name and submitter institution, and each compound design submitted received a unique ID (“PostEra ID”) that appended a unique molecule sequence ID within the submission batch ID. Internally, compound designs, synthesized compounds, and compounds with experimental data were tracked with corresponding records in a CDD Vault (Collaborative Drug Discovery Inc.).

Stereochemistry: While the design platform enabled submitters to register compounds with specific defined or uncertain stereochemistry, compounds were initially synthesized and biochemically assayed as racemates, and if active, chirally separated compounds were registered and assayed separately. Because the absolute stereochemical identity of enantiopure compounds was unknown at time of receipt, assay data was attached to compound records with specified *relative* stereochemistry, rather than absolute stereochemistry. For compounds where sufficient data was available from a variety of sources to propose the absolute stereochemistry (e.g. X-ray data for the compound or a close analogue), the “suspected_SMILES” record was updated, along with an articulated rationale in the “why_suspected_SMILES” field. As a result, caution must be exercised when using data for enantiopure compounds for downstream uses (e.g. whole-dataset machine learning) without verifying the absolute stereochemistry is known with confidence.

Submission analysis: the submitter names were standardised by removing affiliations and expansion of first name abbreviations, the submissions by two users who submitted large batches of compounds in an automated way in contravention of the goal of the project were removed. The word cloud was generated by filtered against 1,000 most common words and removing grammatical inflections and generating an image with an online word cloud generator. The classification of the methodology was done by presence of keywords determined by a simple keyword classifier with manually determined words (circa 100 training, 100 test) wherein ‘dock’, ‘seesar’, ‘vina’, ‘autodock’, ‘screen’, ‘drug-hunter’ were typical of docking, while ‘by-eye’, ‘merg[ing]’, ‘link[ing]’, ‘coupl[ing]’ were typical of hypothesis driven methods. A large fraction could not be accurately classified due to paucity of information. SAScore was calculated with Postera Manifold under the retrosynthesis route.

### 1. Experimental methods

#### 1.1 Protease activity assays

##### 1.1.1 Fluorescence MPro inhibition assay

Compounds were seeded into assay-ready plates (Greiner 384 low volume, cat 784900) using an Echo 555 acoustic dispenser, and DMSO was back-filled for a uniform concentration in assay plates (DMSO concentration maximum 1%) Screening assays were performed in duplicate at 20µM and 50µM. Hits of greater than 50% inhibition at 50µM were confirmed by dose response assays. Dose response assays were performed in 12 point dilutions of 2-fold, typically beginning at 100µM. Highly active compounds were repeated in a similar fashion at lower concentrations beginning at 10µM or 1µM. Reagents for Mpro assay were dispensed into the assay plate in 10µl volumes for a final volume of 20µL.

Final reaction concentrations were 20mM HEPES pH7.3, 1.0mM TCEP, 50mM NaCl, 0.01% Tween-20, 10% glycerol, 5nM Mpro, 375nM fluorogenic peptide substrate ([5-FAM]-AVLQSGFR-[Lys(Dabcyl)]-K-amide). Mpro was pre-incubated for 15 minutes at room temperature with compound before addition of substrate and a further 30 minute incubation. Protease reaction was measured in a BMG Pherastar FS with a 480/520 ex/em filter set. Raw data was mapped and normalized to high (Protease with DMSO) and low (No Protease) controls using Genedata Screener software. Normalized data was then uploaded to CDD Vault (Collaborative Drug Discovery). Dose response curves were generated for IC50 using nonlinear regression with the Levenberg–Marquardt algorithm with minimum inhibition = 0% and maximum inhibition = 100%.

The assay was calibrated at different enzyme concentrations to confirm linearity and response of protease activity, as well as optimization of buffer components for most stable and reproducible assay conditions. Substrate concentration was chosen after titration to minimize saturation of signal in the plate reader while obtaining a satisfactory and robust dynamic range of typically 5-6 fold over control without enzyme. We used low substrate concentrations of the bright FRET peptide to avoid “inner filter effect”(*60*), and to bias towards detection of competitive inhibitors(*61*). As positive control, under our assay condition Nirmatrelvir has IC50 of 2.6nM.

##### 1.1.2 RapidFire MPro inhibition assay

The assay was performed according to the published procedure(*62*). Briefly, compounds were seeded into assay-ready plates (Greiner 384PP, cat# 781280) using an ECHO 650T dispenser and DMSO was back-filled for a uniform concentration in assay plates (DMSO concentration < 1%, final volume = 500 nL.). A 15 µM enzyme stock solution is prepared in 20 mM HEPES, pH 7.5 and 300 mM NaCl, and subsequently diluted to a working solution of 300 nM Mpro in assay buffer (20 mM HEPES, pH 7.5 and 50 mM NaCl) before the addition of 25 µL to each well using a Multidrop Combi (Thermo Scientific). After a quick centrifugation step (1000 rpm, 15 s) the plate is incubated for 15 min at room temperature. The reaction is initiated with the addition of 25 µL of 4 µM 11-mer (TSAVLQSGFRK-NH2, initially custom synthesized by the Schofield group, GLBiochem, used until March 2021), or 10 µM 37-mer (ALNDFSNSGSDVLYQPPQTSITSAVLQSGFRKMAFPS-NH2, GLBiochem, used after March 2021), dissolved in assay buffer. After centrifugation (1000 rpm, 14 s) the reaction is incubated for 10 min (11-mer) or 5 min (37-mer) at room temperature before quenching with 10 % formic acid. The reactions are analysed with MS using RapidFire (RF) 365 high-throughput sampling robot (Agilent) connected to an iFunnel Agilent 6550 accurate mass quadrupole time-of-flight (Q-TOF) mass spectrometer using electrospray. All compounds are triaged by testing the % inhibition at 5 and 50 µM final concentration. Dose response curves uses an 11-point range of 100--0.0017 µM inhibitor concentrations. RapidFire integrator software (Agilent) was used to extract the charged states from the total ion chromatogram data followed by peak integration. For the 11-mer peptide the m/z (+1) charge states of both the substrate (1191.67 Da) and cleaved N-terminal product TSAVLQ (617.34 Da) were used and the 37-mer peptide the m/z (+2) charge states of the substrate (3960.94 Da) and m/z (+1) of the cleaved C-terminal product SGFRKMAFPS (1125.57 Da). Percentage conversion (product peak integral / (product peak integral + substrate peak integral))*100) and percentage inhibitions were calculated and normalised against DMSO control with deduction of any background signal in Microsoft Excel. IC50s were calculated using Levenberg–Marquardt algorithm used to fit a restrained Hill equation to the dose-response data with both GraphPad PRISM and CDD.

#### 1.2 High throughput X-ray crystallography

Purified protein(*20*) at 24 mg/ml in 20 mM HEPES pH 7.5, 50 mM NaCl buffer was diluted to 12 mg/ml with 20 mM HEPES pH 7.5, 50 mM NaCl before performing crystallization using the sitting-drop vapour diffusion method with a reservoir solution containing 11% PEG 4 K, 5% DMSO, 0.1 M MES pH 6.5. Crystals of Mpro in the monoclinic crystal form (C2), with a single monomer in the asymmetric unit, were grown with drop ratios of 0.15 µl protein, 0.3 µl reservoir solution and 0.05 µl seeds prepared from previously produced crystals of the same crystal form(*20*). Crystals in the orthorhombic crystal form (P2_1_2_1_2_1_), with the Mpro dimer present in the asymmetric unit, were grown with drop ratios of 0.15 µl protein, 0.15 µl reservoir solution and 0.05 µl seeds prepared from crystals of an immature Mpro mutant in the same crystal form(*63*).

Compounds were soaked into crystals by adding compound stock solutions directly to the crystallisation drops using an ECHO liquid handler. In brief, 40-90 nl of DMSO solutions (between 20 and 100 mM) were transferred directly to crystallisation drops using giving a final compound concentration of 2-20 mM and DMSO concentration of 10-20%. Drops were incubated at room temperature for approx. 1-3 h prior to mounting and flash cooling in liquid nitrogen without the addition of further cryoprotectant.

Data was collected at Diamond Light Source on the beamline I04-1 at 100 K and processed with the fully automated pipelines at Diamond(*64–66*), which include XDS(*67*), xia2(*68*), autoPROC(*69*) and DIALS(*64*). Further analysis was performed using XChemExplorer(*70*) with electron density maps generated using DIMPLE (http://ccp4.github.io/dimple/). Ligand-binding events were identified using PanDDA(*71*) (https://github.com/ConorFWild/pandda) and ligands were manually modelled into PanDDA-calculated event maps or electron density maps using Coot(*72*). Ligand restraints were calculated with ACEDRG(*73*) or GRADE (grade v. 1.2.19 (Global Phasing Ltd., Cambridge, United Kingdom, 2010)) and structures refined with Buster (Buster v. 2.10.13 (Cambridge, United Kingdom, 2017)). Models and quality annotations were reviewed using XChemReview(*74*), Buster-Report (Buster v. 2.10.13 (Cambridge, United Kingdom, 2017)) and Mogul(*75*, *76*).

Coordinates, structure factors and PanDDA event maps for all data sets are available on Fragalysis (https://fragalysis.diamond.ac.uk/viewer/react/preview/target/Mpro).

#### 1.3 Viral screening assays

A variety of antiviral replication assays were performed in collaborating laboratories, including cytopathic effect (CPE) inhibition assays at the IIBR, Israel, and Katholieke Universiteit Leuven, RT-qPCR for viral RNA at Radboud University Medical Center, Netherlands, immunofluorescence assays at University of Nebraska Medical centre, USA, and plaque assays and focus forming unit assays at University of Oxford, UK.

##### 1.3.1 Antiviral Cytopathic Effect Assay, VeroE6 (IIBR, Ness-Ziona, Israel)

SARS-CoV-2 (GISAID accession EPI_ISL_406862) was kindly provided by Bundeswehr Institute of Microbiology, Munich, Germany. Virus stocks were propagated (4 passages) and tittered on Vero E6 cells. Handling and working with SARS-CoV-2 virus was conducted in a BSL3 facility in accordance with the biosafety guidelines of the Israel Institute for Biological Research (IIBR). Vero E6 were plated in 96-well plates and treated with compounds in medium containing 2 % fetal bovine serum. The assay plates containing compound dilutions and cells were incubated for 1 hour at 37℃ temperature prior to adding Multiplicity of infection (MOI) 0.01 of viruses. Viruses were added to the entire plate, including virus control wells that did not contain test compound and Remdesivir drug used as positive control. After 72h incubation viral cytopathic effect (CPE) inhibition assay was measured with XTT reagent. Three replicate plates were used.

##### 1.3.2 Antiviral Immunoflourescence assay, VeroE6 (Pathology and Microbiology, University of Nebraska Medical Centre, USA, St Patrick Reid)

Vero E6 cells were pretreated with 20 uM of the Moonshot compounds for around 2h. Cells were then infected with SARS-CoV-2 at a MOI of 0.1 for 24h. Virus infection was terminated by 4 % PFA fixation. Cells were stained using a Rabbit SARS-CoV-2 antibody (Sino Biological 40150-R007) as a primary antibody, and Alexa-488, Hoechst and Cell Mask (Thermo Fisher) as a secondary antibody. Images were collected on the Operetta system imaging system, and analysed using the Harmony software.

##### 1.3.3 Antiviral Focus Forming Unit Assay, Calu-3 (University of Oxford, UK)

***Cell culture***. The African green monkey Vero E6 cell line (ATCC CRL-1586) was cultured in Dulbecco’s modified Eagle medium (DMEM) with Glutamax supplemented with 100 µg/mL streptomycin, 100 U/mL penicillin, and 10 % heat-inactivated fetal calf serum (FCS). The human lung cancer cell line Calu-3 (Anderson Ryan, Department of Oncology, Medical Science Division, University of Oxford) was cultured in a 1:1 mixture of DMEM with Glutamax and Ham’s F-12 medium supplemented with 100 µg/mL streptomycin, 100 U/mL penicillin, and 10% heat-inactivated FCS. All cells were maintained as mycoplasma free, with regular verifications by polymerase chain reaction (PCR).

**Virus propagation**. SARS-CoV-2 England/2/2020 was provided at passage 1 from Public Health England, Collindale. Passage 2 submaster and passage 3 working stocks were produced by infecting Vero E6 cells at a multiplicity of infection of 0.01 in virus propagation medium (DMEM with Glutamax supplemented with 2 % FCS) and incubating until cytopathic effect was visible. The cell supernatant was then centrifuged at 500 g for 5 minutes, aliquoted and stored at −80°C. The titre of viral stocks was determined by plaque assay. All subsequent assays were performed using a passage 3 stock.

**Cell viability**. Cell viability was was measured using the CellTiter 96 R AQueous One Solution Cell Proliferation MTA (3-(4,5-dimethylthiazol-2-yl)-5-(3-carboxymethoxyphenyl)-2-(4-sulfophenyl)-2H - 15 tetrazolium, inner salt) Assay (Promega) according to the manufacturer’s instruction after treatment with compound. Briefly, Calu 3 cells were treated with compounds in quadruplicate for 3 days. Wells with 200 µL growth medium with and without cells were included as controls in quadruplicate. After the incubation, 100 µL of growth medium was removed and 20 µL of MTS reagent was added to the remaining medium in each well. After a further one to two hour incubation, the absorbance at 490 nm was measured on a Molecular Devices SpectraMax M5 microplate reader.

***Antiviral assays.*** For Focus forming unit assays, a SARS-CoV-2 Microneutralization assay from the W James lab (Dunn School of Pathology, University of Oxford) was adapted for use as a FFU assay. Briefly, 3 half log dilutions of each supernatant to be analyzed were prepared in virus propagation medium. 20µL of each dilution was inoculated into wells of a 96-well plate in quadruplicate followed by 100 μL Vero E6 cells at 4.5 x 10^5 cells/mL in virus propagation medium. The plates were incubated for 2 hours prior to the addition of 100 μL of 1.8 % CMC overlay, and then incubated for a further 24 hours. After 24 hours the overlay was carefully removed and the cells washed once with PBS before fixing with 50µL of 4 % paraformaldehyde, after 30 minutes the paraformaldehyde was removed and replaced with 100µL of 1 % ethanolamine in PBS. The cells were permeabilized by replacing the ethanolamine with 2 % Triton X100 in PBS and incubating at 37°C for 30 minutes. The plates were then washed 3 times with wash buffer (0.1 % Tween 20 in PBS) inverted and gently tapped onto tissue to dry before the addition of 50 μl of EY2A anti-N human mAb (Arthur Huang (Taiwan)/Alain Townsend (Weatherall Institute of Molecular Medicine, University of Oxford)) at 10 pmol in wash buffer. The plates were rocked at room temperature for 1 hour, washed and incubated with 100μl of secondary antibody Anti-Human IgG (Fc-specific)-peroxidase-conjugate produced in Goat diluted 1:5000 at room temperature for 1 hour. 50µL of TrueBlue peroxidase substrate was added to the wells and incubated at RT for 10 min on the rocker, after 10 minutes the substrate was removed and the plates washed with ddH2O for 10 minutes. The H2O was removed and the plates allowed to air dry. The foci were then counted using an ELISPOT classic reader system (AID GmbH).

##### 1.3.4 Antiviral qPCR assay, VeroE6 and kidney organoids (Radboud University Medical Center, Nijmegen, Netherlands)

***Cell culture*** African green monkey Vero E6 kidney cells (ATCC CRL-1586) and Vero FM kidney cells (ATCC CCL-81) were cultured in Dulbecco’s modified Eagle medium (DMEM) with 4.5 g/L glucose and L-glutamine (Gibco), supplemented with 10% fetal calf serum (FCS, Sigma Aldrich), 100 µg/ml streptomycin and 100 U/ml penicillin (Gibco). Cells were maintained at 37 °C with 5% CO_2_. Human induced pluripotent stem cell-derived kidney organoids were prepared as previously described(*52*).

***Virus propagation.*** SARS-CoV-2 (isolate BetaCoV/Munich/BavPat1/2020) was kindly provided by Prof. C. Drosten (Charité-Universitätsmedizin Berlin, Institute of Virology, Berlin, Germany) and was initially cultured in Vero E6 cells up to three passages in the laboratory of Prof. Bart Haagmans (Viroscience Department, Erasmus Medical Center, Rotterdam, The Netherlands). Vero FM cells were infected with passage 3 stock at an MOI of 0.01 in infection medium (DMEM containing L-glutamine, 2% FCS, 20 mM HEPES buffer, 100 µg/ml streptomycin and 100 U/ml penicillin). Cell culture supernatant containing virus was harvested at 48 hours post-infection (hpi), centrifuged to remove cellular debris, filtered using a 0.2 μm syringe filter (Whatman), and stored in 100 μl aliquots at −80°C.

***Virus titration.*** Vero E6 cells were seeded in 12-well plates at a density of 500,000 cells/well. Cell culture medium was discarded at 24 h post-seeding, cells were washed twice with PBS and infected with 10-fold dilutions of the virus stock in unsupplemented DMEM. At 1 hpi, cells were washed with PBS and replaced with overlay medium, consisting of Minimum Essential medium (Gibco), 2% FCS, 20 mM HEPES buffer, 100 μg/ml streptomycin, 100 U/ml penicillin and 0.75% carboxymethyl cellulose (Sigma Aldrich). At 72 hpi, the overlay medium was discarded, cells were washed with PBS and stained with 0.25% crystal violet solution containing 4% formaldehyde for 30 minutes. Afterwards, staining solution was discarded and plates were washed with PBS, dried and plaques were counted.

***Antiviral assay.*** Vero E6 cells were seeded onto 24-well plates at a density of 150,000 cells/well. At 24 h post-seeding, cell culture medium was discarded, cells were washed twice with PBS and infected with SARS-CoV-2 at an MOI of 0.01 in the presence of six concentrations of the inhibitors (25 μM – 0.06 μM). At 1 hpi, the inoculum was discarded, cells were washed with PBS, and infection medium containing the same concentration of the inhibitors was added to the wells. SARS-CoV-2 infection in the presence of 0.1% DMSO was used as a negative control. At 24 hpi, 100 μl of the cell culture supernatant was added to RNA-Solv reagent (Omega Bio-Tek) and RNA was isolated and precipitated in the presence of glycogen according to manufacturer’s instructions. TaqMan Reverse Transcription reagent and random hexamers (Applied Biosystems) were used for cDNA synthesis. Semi-quantitative real-time PCR was performed using GoTaq qPCR (Promega) BRYT Green Dye-based kit using primers targeting the SARS-CoV-2 E protein gene(*77*) (Forward primer, 5′-ACAGGTACGTTAATAGTTAATAGCGT-3’; Reverse primer, 5′-ACAGGTACGTTAATAGTTAATAGCGT-3’). A standard curve of a plasmid containing the E gene qPCR amplicon was used to convert Ct values relative genome copy numbers. For viability assays, Vero E6 cells were seeded in 96-well white-bottom culture plates (Perkin Elmer) at a density of 30,000 cells per well. At 24 h post-seeding, cells were treated with the same concentrations of compounds as used for the antiviral assay. Cells treated with 0.1 % DMSO were used as a negative control. At 24 h post-treatment, cell viability was assessed using the Cell Titer Glo 2.0 kit (Promega) using the Victor Multilabel Plate Reader (Perkin Elmer) to measure luminescence signal.

***Antiviral assays in organoids***. Human iPSC-derived kidney organoids cultured in transwell filters (Corning) were infected with SARS-CoV-2 in the presence of 1 and 10 µM of MAT-POS-932d1078-3, PET-UNK-29afea89-2 or 0.1% DMSO using an MOI of 1.0 in Essential 6 medium (Gibco) at 37°C and 5 % CO_2_, exposing the cells both basolaterally and apically to the inoculum. After 24 h, medium containing the inoculum was removed and fresh Essential 6 medium containing the same concentration of inhibitor was added to the basolateral compartment and cells were cultured for an additional 24 h. At 48 hpi, organoids were washed in PBS, and the apical surface was exposed to Essential 6 medium for 10 min at 37°C, which was collected and used for viral titration. Individual organoids were harvested for RNA isolation using the PureLink RNA mini kit (Thermo Fisher) according to manufacturer’s instructions. Viral RNA copies were analyzed by RT-qPCR on the SARS-CoV E gene, as described previously(*78*).

##### 1.3.5 High-content SARS-CoV-2 antiviral screening assay, Hela-ACE2 (Takeda via Calibr/TSRI)

***SARS-CoV-2/HeLa-ACE2 high-content screening assay.*** Compounds are acoustically transferred into 384-well µclear-bottom plates (Greiner, Part. No. 781090-2B) and HeLa-ACE2 cells are seeded in the plates in 2% FBS at a density of 1.0×10^3^ cells per well. Plated cells are transported to the BSL3 facility where SARS-CoV-2 (strain USA-WA1/2020 propagated in Vero E6 cells) diluted in assay media is added to achieve ∼30 – 50% infected cells. Plates are incubated for 24 h at 34℃ 5% CO_2_, and then fixed with 8% formaldehyde. Fixed cells are stained with human polyclonal sera as the primary antibody, goat anti-human H+L conjugated Alexa 488 (Thermo Fisher Scientific A11013) as the secondary antibody, and antifade-46-diamidino-2-phenylindole (DAPI; Thermo Fisher Scientific D1306) to stain DNA, with PBS 0.05% Tween 20 washes in between fixation and subsequent primary and secondary antibody staining. Plates are imaged using the ImageXpress Micro Confocal High-Content Imaging System (Molecular Devices) with a 10× objective, with 4 fields imaged per well. Images are analyzed using the Multi-Wavelength Cell Scoring Application Module (MetaXpress), with DAPI staining identifying the host-cell nuclei (the total number of cells in the images) and the SARS-CoV-2 immunofluorescence signal leading to identification of infected cells.

***Uninfected host cell cytotoxicity counter screen.*** Compounds are acoustically transferred into 1,536-well plates (Corning No. 9006BC). HeLa-ACE2 cells are maintained as described for the infection assay and seeded in the assay-ready plates at 400 cells/well in DMEM with 2% FBS. Plates are incubated for 24 hours at 37℃ 5% CO_2_. To assess cell viability, 2 mL of 50% Cell-Titer Glo (Promega No G7573) diluted in water is added to the cells and luminescence measured on an EnVision Plate Reader (Perkin Elmer).

***Data analysis.*** Primary in vitro screen and the host cell cytotoxicity counter screen data are uploaded to Genedata Screener, Version 16.0. Data are normalized to neutral (DMSO) minus inhibitor controls (2.5 µM remdesivir for antiviral effect and 10 µM puromycin for infected host cell toxicity). For the uninfected host cell cytotoxicity counter screen 40 µM puromycin (Sigma) is used as the positive control. For dose response experiments compounds are tested in technical triplicates on different assay plates and dose curves are fitted with the four parameter Hill Equation.

##### 1.3.6 Cytopathic Effect Assay, hACE2-TMPRSS2 cells (Katholieke Universiteit Leuven)

***Virus isolation and virus stocks*** All virus-related work was conducted in the high-containment BSL3 facilities of the KU Leuven Rega Institute (3CAPS) under licenses AMV 30112018 SBB 219 2018 0892 and AMV 23102017 SBB 219 2017 0589 according to institutional guidelines. The Severe Acute Respiratory Syndrome Coronavirus 2 (SARS-CoV-2) strain used for this study was the Alpha variant of Concern (derived from hCoV-19/Belgium/rega-12211513/2020; EPI_ISL_791333, 2020-12-21). Virus sample was originally isolated in-house from nasopharyngeal swabs taken from travellers returning to Belgium (baseline surveillance) and were subjected to sequencing on a MinION platform (Oxford Nanopore) directly from the nasopharyngeal swabs. Virus stocks were then grown on Vero E6 cells in (DMEM 2% FBS medium) and passaged one time on A549-ACE2TMPRSS2 cells. Median tissue culture infectious doses (TCID50) was defined by end-point titration.

***A549-ACE2-TMPRSS2 assay*** A549-Dual™ hACE2-TMPRSS2 cells obtained by Invitrogen (Cat. a549d-cov2r) were cultured in DMEM 10% FCS (Hyclone) supplemented with 10 µg/ml blasticidin (Invivogen, ant-bl-05), 100 µg/ml hygromycin (Invivogen, ant-hg-1), 0.5 µg/ml puromycin (Invivogen, ant-pr-1) and 100 µg/ml zeocin (Invivogen, ant-zn-05). For antiviral assay, cells were seeded in assay medium (DMEM 2%) at a density of 15,000 cells/well. One day after, compounds were serially diluted in assay medium (DMEM supplemented with 2% v/v FCS) and cells were infected with their respective SARS-CoV-2 strain at a MOI of approximately 0.003 TCID50/ml. On day 4 pi., differences in cell viability caused by virus-induced CPE or by compound-specific side effects were analyzed using MTS as described previously(*79*). Cytotoxic effects caused by compound treatment alone were monitored in parallel plates containing mock-infected cells.

##### 1.3.6 Immunofluorescence SARS-CoV-2 antiviral screening assay, Hela-ACE2 (Mount Sinai)

Assessment of cross-reactivity against SARS-CoV-2 variant strains and cytotoxicity assays were performed as previously described(*80*). In brief, two thousand HeLa-ACE2 cells (BPS Bioscience) were seeded into 96-well plates in DMEM (10% FBS) and incubated for 24 hours at 37°C, 5% CO2. Two hours before infection, the medium was replaced with 100 μL of DMEM (2% FBS) containing the compound of interest at concentrations 50% greater than those indicated, including a DMSO control. Plates were then transferred into the BSL3 facility and 100 PFU (MOI = 0.025) was added in 50 μL of DMEM (2% FBS), bringing the final compound concentration to those indicated. Plates were then incubated for 48 hours at 37°C. After infection, supernatants were removed and cells were fixed with 4% formaldehyde for 24 hours prior to being removed from the BSL3 facility. The cells were then immunostained for the viral N protein (an inhouse mAb 1C7, provided by Dr. Thomas Moran) with a DAPI counterstain. Infected cells (488 nm) and total cells (DAPI) were quantified using the Celigo (Nexcelcom) imaging cytometer. Infectivity was measured by the accumulation of viral N protein (fluorescence accumulation). Percent infection was quantified as ((Infected cells/Total cells) - Background) *100 and the DMSO control was then set to 100% infection for analysis. Data was fit using nonlinear regression and IC50s for each experiment were determined using GraphPad Prism version 8.0.0 (San Diego, CA). Cytotoxicity was also performed using the MTT assay (Roche), according to the manufacturer’s instructions. Cytotoxicity was performed in uninfected cells with same compound dilutions and concurrent with viral replication assay. All assays were performed in biologically independent triplicates.

### 2. Computational Methods

#### 2.1 Synthetic route planning

We employ an approach based on the Molecular Transformer technology(*32*). Our algorithm uses natural language processing to predict the outcomes of chemical reactions and design retrosynthetic routes starting from commercially available building blocks. This proprietary platform is provided free of charge by PostEra Inc (http://postera.ai). Additionally, Manifold (https://postera.ai/manifold) was built by PostEra Inc. during the project to search the entire space of purchasable molecules, and automatically find the optimal building blocks.

#### 2.2 Alchemical free energy calculations

Large-scale alchemical free energy calculations were conducted in batches (“Sprints”) in which each set of calculations aimed to prioritize compounds that could be produced from a common synthetic intermediate using Enamine’s extensive building block library, resulting in synthetic libraries of hundreds to tens of thousands. Virtual synthetic libraries were organized into a star map, where all transformations were made with respect to a single reference X-ray structure and compound with experimentally measured bioactivity. X-ray structures were prepared using the OpenEye Toolkit SpruceTK with manually controlled protonation states for the key His41:Cys145 catalytic dyad (variously using zwitterionic or uncharged states) and His163 in P1 (which interacts with the 3-aminopyridine or isoquinoline nitrogen in our primary lead series). As the most relevant protonation states were uncertain, when computational resources afforded, calculations were carried out using multiple protonation state variants (His41:Cys145 either neutral or zwitterionic; His163 neutral or protonated) and the most predictive model on available retrospective data for that scaffold selected for nominating prospective predictions for that batch. Initial poses of target compounds were were generated via constrained conformer enumeration to identify minimally-clashing poses using Omega (from the OpenEye Toolkit) using a strategy that closely follows an exercise described in a blog post by Pat Walters (http://practicalcheminformatics.blogspot.com/2020/03/building-on-fragments-from-diamondxchem_30.html). Alchemical free energy calculations were then prepared using the open source perses relative alchemical free energy toolkit(*40*) (https://github.com/choderalab/perses), and nonequilibrium switching alchemical free energy calculations(*81*) were run on Folding@home using the OpenMM compute core(*43*). Nonequilibrium switching calculations used 1 ns nonequilibrium alchemical trajectories, where most calculations were performed with 1 fs timesteps without constraints to hydrogen due to technical limitations that have been resolved in calculations employing OpenMM 7.5.1 and later. We used the Open Force Field Initiative OpenFF “Parsley” small molecule force fields(*39*) (multiple generations between 1.1.1 and 1.3.1 were released and used as the project evolved) and the AMBER14SB protein force field(*82*) with recommended ion parameters(*83*, *84*), and TIP3P water(*85*). As many assayed compounds as possible were included in each batch of transformations to enable continual retrospective assessment and to leverage existing measured affinities in extrapolating predicted affinities. Analysis of free energy calculations used the maximum likelihood estimator [https://doi.org/10.1021/acs.jcim.9b00528] to reconstruct the optimal predicted absolute free energy (and hence pIC50) estimate from available experimental measurements. Calculations were analyzed using the fah-xchem dashboard (https://github.com/choderalab/fah-xchem) using the Bennett acceptance ratio(*86*, *87*) (https://threeplusone.com/pubs/gecthesis) and posted online in real time for the medicinal chemistry team to consult in making decisions about which compounds to prioritize.

We note that our primary aim was computing estimates of relative binding free energies for large alchemical transformations using abundant computing resources (which exceeded 1 exaFLOP/s(*38*)) rather than aggressive optimization of the cost/transformation. Batches of transformations used between 100-200 parallel 4 ns nonequilibrium cycles per transformation, selected based on the number of atoms modified in the transformation, resulting in 100-200 ns/transformation in aggregate. A Tesla V100 achieves ∼200 ns/day for our solvated Mpro complex, meaning ∼2-4 GPU-days/transformation was consumed on a V100 equivalent GPU. To give typical scales, **Figure 2C Panel 1** ran 6319 transformations of 140 cycles, resulting in ∼3.5 ms of simulation time or ∼424K GPU-hours; **Figure 2C Panel 2** ran 5077 transformations of ∼200 cycles, resulting in ∼4 ms simulation time, or ∼480K GPU-hours; **Figure 2C Panel 3** ran 686 transformations of ∼200 cycles, resulting in ∼548 µs of simulation time, or ∼66K GPU-hours.

Scripts for setting up and analysing the perses alchemical free energy calculations on Folding@home, as well as an index of computed datasets and dashboards are available at https://github.com/foldingathome/covid-moonshot

Code used for generating the COVID Moonshot alchemical free energy calculation web dashboards is available here: https://github.com/choderalab/fah-xchem Retrospective calculations for transformations in the main synthetic series shown in **Figure 5A** were performed with an early release of perses 0.10.2 constructed as a simplified example that anyone can run to illustrate how these calculations work on standard GPU workstations, and use standard alchemical replica exchange protocols of 5 ns/replica (which just take a few hours on standard workstations, as opposed to the expensive nonequilibrium protocols used in the Sprints). Input scripts for this calculation are available in the perses distribution under ‘examples/moonshot-mainseries/’ [https://github.com/choderalab/perses/tree/main/examples/moonshot-mainseries].

#### 2.3 Structural flexibility and interactions analysis

Protein-ligand interactions are the driving forces for molecular recognition. In this work, the *PLIPify* repo (https://github.com/volkamerlab/plipify) is used to detect shared interaction hot spots within the different MPro structures. *PLIPify* is a python wrapper built on top of PLIP(*88*), a tool that enables automatic generation of protein-ligand interaction profiles for single complexes, to allow combining these profiles for multiple structures.

To generate the hotspots (depicted in **Figure 3A**), the fragalysis data was downloaded (as of July 2021, https://fragalysis.diamond.ac.uk/api/targets/?format=json&title=Mpro). The respective pre-aligned complex structures were further investigated (found under data/{target}/aligned/{crystal_name}/{crystal_name}_bound.pdb). Only one chain per structure is kept, and the structures are protonated using Amber’s *reduce* function. *PLIPify* is invoked and structures are excluded from further analysis if they do not contain exactly one binding site (i.e. PLIP detects either zero or more than 1 binding sites), the sequence contains gaps (‘-’) or the sequence length differs more than a standard deviation from the average length across all investigated structures.

This resulted in a final set of 367 complex structures, used to generate the interaction fingerprints. Note for this study, only hbond-donor, hbond-acceptor, salt bridge, hydrophobic, pi-stacking, and halogen interactions are inspected. Additional code was added to *PLIPify* to split the hbond-donor and hbond-acceptor interactions into backbone and sidechain interactions (https://github.com/volkamerlab/plipify/pull/18). Interacting residues are only included if the summed interaction count per residue over all investigated structures is greater than five. Careful examination of examples of the interactions led us to filter out the S144 interactions from the final report as none of the interactions were convincing (24 hbond-don-bb, 168 hbond-don-sc, and 4 hbond-acc-sc interactions). The resulting structural depiction (**Figure 3A**) were generated using pymol, and structure Mpro-P1788_0A_bound_chainA (protonated) is displayed (scripts available at https://github.com/volkamerlab/plipify/tree/master/projects/01/fragalysis.ipynb). Finally, structures containing compounds exhibiting some of the major interactions identified were used to generate the figures in **Figure 3B**.

### 4. Chemical methods

#### 4.1 HTC library synthesis

##### 4.1.1 Chan-Lam reaction

The arylamine library was made by reacting the boronic acid (**Fig S7D**), under the optimized reaction conditions (1 eq. amine; 0.2 eq. CuI; 0.8 eq. DMAP; 2 eq. Hex_3_N; DMSO; under air; RT; 2 days) with 296 amines (200 aromatic, 48 primary, and 48 secondary aliphatic amines; **Data S2**). For library production, we used Echo LDV plates and an Echo 555 acoustic dispenser for liquid handling. After the allotted reaction time, plate copies were made after diluting the reaction mixture with 4.6 μL DMSO and transferring 1 μL of the obtained solution to an 384-well plate, for either biochemical assay or yield estimation.

##### 4.1.2 Amide coupling

The amide library was made by reacting the carboxylic acid (**Fig S8E**) under the optimized reaction conditions (2 eq. amine; 2 eq. EDC; 2 eq. HOAt; 5 eq. DIPEA; DMSO; RT; 24h) with 300 amines (202 aromatics, 49 primary, and 49 secondary aliphatic amines Data S2). For library production, we used Echo LDV plates and an Echo 555 acoustic dispenser for liquid handling. Plate copies were made after diluting the reaction mixture with 4 μL DMSO. For yield estimation, 1 μL of the diluted library was transferred to an LC/MS-ready 384-well plate, followed by dilution with 20% ACN in water to the final volume of 50 μL. The desired product was identified in 60% of wells.

#### 4.2 General compounds synthesis and characterization

All compounds were directly purchased from Enamine Inc., following Enamine’s standard QC for compound collections. In addition, in Supplementary Chemistry we discuss the synthesis procedure, as well as LCMS and 1H NMR characterisation of compounds which were discussed in the manuscript with associated bioactivity data.

All COVID Moonshot compounds are publicly available as a screening collection that can be ordered in bulk or as singleton through Enamine. The compound identifiers of the COVID Moonshot collection are in the Supplementary Data, together with Enamine’s internal QC data comprising LCMS spectra for all compounds and NMR spectra for selected compounds.

## Acknowledgements

The COVID Moonshot acknowledges funding by the Wellcome Trust on behalf of the Covid-19 Therapeutics Accelerator. The COVID Moonshot project is particularly grateful to UCB Pharma Ltd and UCB S.A for the support from their Medicinal and Computational Chemistry groups, to Novartis Institute for Biomedical Research for generous in-kind ADME and PK contributions, Takeda for in-kind contribution of antiviral assays/pan-corona biochemical assays, and Nanosyn for protease panel assays. We thank CDD Vault and OpenEye Scientific for their in-kind contribution allowing the consortium to use their software. We also thank the numerous volunteers that contributed compound designs to the COVID Moonshot and to the citizen scientist volunteers of Folding@home for donating their computing resources, and Amazon Web Services for key support of Folding@home infrastructure.

## Funding Acknowledgments and Financial Disclosures

Funding acknowledgements and Disclosures statements for each author are listed in the consortium spreadsheet (**Data S7**). This research was funded in part, by the Wellcome Trust [Grant number 209407/Z/17/Z]. For the purpose of open access, the author has applied a CC-BY public copyright license to any Author Accepted Manuscript version arising from this submission.

## Data availability

- **All compound designs, datasets, and X-ray structures can be browsed on the COVID Moonshot website**: https://postera.ai/covid; The compound submissions and experimental data are available via GitHub https://github.com/postera-ai/COVID_moonshot_submissions and the bioactivity data can be interactively browsed https://covid.postera.ai/covid/activity_data; All of the data is also available in a permanent archive(*89*)
- **Alchemical free energy calculations code and datasets are indexed on GitHub and stored in a permanent archive**(*90*) https://github.com/foldingathome/covid-moonshot
- **All X-ray structures are available for interactive viewing and comparison or bulk download via Fragalysis:** https://fragalysis.diamond.ac.uk/viewer/react/preview/target/Mpro; Structures were deposited to the PDB (**Data S4**) and are also available in a permanent archive.(*91*)

## Resource availability

- **Synthesized compounds**: We have made all compounds assayed here available from the current Enamine catalog, and readily available for purchase from Enamine (and other suppliers) via the Manifold platform accessible for each compound page on the COVID Moonshot website: https://covid.postera.ai/covid/activity_data

## Supplementary Materials

Materials and Methods

Supplementary Chemistry

Figs. S1 to S18

Data S1-S7

References 93-94

