## Supplemental Materials for "Open Science Discovery of Potent Non-Covalent SARS-CoV-2 Main Protease Inhibitors"

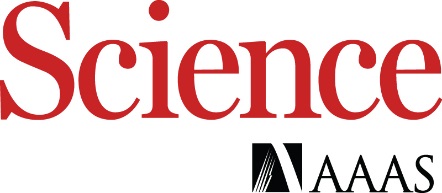

Supplementary Materials for

**Open Science Discovery of Potent Non-Covalent SARS-CoV-2 Main Protease Inhibitors**

Melissa L. Boby^1,2,3,^**^*^,** Daren Fearon^4,5,^**^*^,** Matteo Ferla^6,^**^*^,** Mihajlo Filep^7,*^, Lizbé Koekemoer^8,*^, Matthew C. Robinson^9,^**^*^,** [The COVID Moonshot Consortium](https://docs.google.com/spreadsheets/d/1OnPzj5IvYtD5uu6tqmlhvOX47__WgN1VrbzMH_kqns0/edit?pli=1#gid=0), John D. Chodera^3,#^, Alpha A Lee^9,#^, Nir London^7,#^, Annette von Delft^6,8,10,#^, Frank von Delft^4,5,8,10,11,#^

### ^1^ Pharmacology Graduate Program, Weill Cornell Graduate School of Medical Sciences, New York, NY 10065, USA

### ^2^ Program in Chemical Biology, Sloan Kettering Institute, Memorial Sloan Kettering Cancer Center, New York, NY 10065, USA

### ^3^ Program in Computational and Systems Biology, Sloan Kettering Institute, Memorial Sloan Kettering Cancer Center, New York, NY 10065, USA

^4^ Diamond Light Source Ltd., Harwell Science and Innovation Campus, Didcot, OX11 0QX, UK

^5^ Research Complex at Harwell, Harwell Science and Innovation Campus, Didcot OX11 0FA, United Kingdom

^6^ Oxford Biomedical Research Centre, National Institute for Health Research, University of Oxford, Oxford, UK

^7^ Department of Chemical and Structural Biology, The Weizmann Institute of Science, Rehovot, Israel, 7610001

^8^ Centre for Medicines Discovery, Nuffield Department of Medicine, University of Oxford, Oxford, UK

^9^ PostEra Inc., 1 Broadway, 14th Floor,Cambridge, MA 02142, USA

### ^10^ Structural Genomics Consortium, Nuffield Department of Medicine, University of Oxford, Oxford, UK

^11^ Department of Biochemistry, University of Johannesburg, Auckland Park, Johannesburg 2006, South Africa

### ^*^ Equal contribution ; Authors are ordered alphabetically.

### ^#^ For correspondence: (John D. Chodera); (Alpha Lee); (Nir London); (Annette von Delft); (Frank von Delft)

**The PDF file includes:**

Supplementary Chemistry

Figs. S1 to S18

Table S1

**Other Supplementary Materials for this manuscript include the following:**

Data S1 to S7

Supplementary Chemistry

**Abbreviations Used**

EDC, 1-ethyl-3-(3-dimethylaminopropyl)carbodiimide;

TEA, triethylamine;

CDI, 1,1′-carbonyldiimidazole;

DMSO, dimethyl sulfoxide;

DIPEA, N-ethyl-N-isopropylpropan-2-amine;

DCM, dichloromethane;

HOAc, acetic acid;

MeOH, methanol;

DMSO, dimethyl sulfoxide;

HPLC, high-performance liquid chromatography;

LCMS, liquid chromatography-mass spectrometry;

min, minute(s);

h, hour(s);

HOAt, 1-hydroxy-7-azabenzotriazole;

HATU, *N*-[(dimethylamino)-1*H*-1,2,3-triazolo-[4,5-*b*]pyridin-1-ylmethylene]-*N*-methylmethanaminium hexafluorophosphate *N*-oxide;

IPA, isopropanol;

RT, retention time;

equiv., equivalent;

TFA, trifluoroacetic acid;

**General procedure**

**2-(3-Chlorophenyl)-N-(isoquinolin-4-yl)acetamide (Z1530724813)**

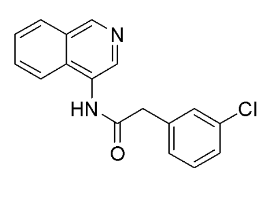

Isoquinolin-4-amine hydrochloride (61.6 mg, 342.14 mmol), 2-(3-chlorophenyl)acetic acid (64.0 mg, 376.45 mmol), EDC (64.1 mg, 413.18 mmol), TEA (41.4 mg, 409.41 mmol), and HOAt (50.82 mmol, 410.12 mmol) were mixed in anhydrous DMSO (0.5 mL). The reaction mixture was sealed and kept at 25 °C for 18 h. After that, the mixture was evaporated under reduced pressure, and the residue was dissolved in DMSO (0.6 mL). The solution was filtered, analyzed by LCMS, and then purified by HPLC to afford 2-(3-chlorophenyl)-N-(isoquinolin-4-yl)acetamide (32.0 mg, 31.62%) as a yellow solid.

**^1^H NMR (500 MHz, DMSO-*d_6_*)** δ (ppm) 10.26 (s, 1H), 9.14 (s, 1H), 8.68 (s, 1H), 8.14 (d, *J* = 8.1 Hz, 1H), 8.05 (d, *J* = 8.5 Hz, 1H), 7.82 (dd, *J* = 8.4, 6.9 Hz, 1H), 7.70 (t, *J* = 7.5, 7.5 Hz, 1H), 7.48 (s, 1H), 7.42-7.36 (m, 2H), 7.36-7.30 (m, 1H), 3.87 (s, 2H).

**MS (ESI+)** m/z calculated for C_17_H_14_ClN_2_O^+^ ([M+H]^+^) 297.1, found: 297.0.

**HPLC condition:**

Column: Chromatorex 18 SMB100-5T 100x19 mm)

Mobile phase: 40:40:90% at 0:1:5 min, H_2_O/MeOH

Flow rate: 30 mL/min

**2-(3-Chlorophenyl)-N-(4-methylpyridin-3-yl)acetamide (Z1129289650)**

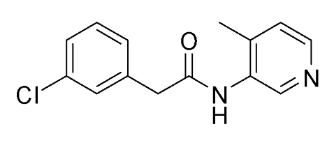

4-Methylpyridin-3-amine (40.9 mg, 378.46 mmol), 2-(3-chlorophenyl)acetic acid (71.0 mg, 417.62 mmol), EDC (71.1 mg, 458.30 mmol), and HOAt (53.69 mg, 438.50 mmol) were mixed in anhydrous DMSO (0.5 mL). The reaction mixture was sealed and kept at 25 °C for 18 h. After that, the mixture was evaporated under reduced pressure; and the residue was dissolved in DMSO (0.7 mL). The solution was filtered, analyzed by LCMS, and then purified by HPLC ( eluting: 40-40-90% 0-1-5 min H_2_O/MeOH, flow: 30 mL/min (loading pump 4 mL/min methanol); column: Chromatorex 18 SMB100-5T 100x19 mm) to afford 2-(3-chlorophenyl)-N-(4-methylpyridin-3-yl)acetamide (62.6 mg, 63.46%) as a yellow solid.

**^1^H NMR (500 MHz, DMSO-*d_6_*)** δ (ppm) 9.76 (s, 1H), 8.47 (s, 1H), 8.22 (d, *J* = 5.0 Hz, 1H), 7.44-7.39 (m, 1H), 7.39-7.34 (m, 1H), 7.34-7.28 (m, 2H), 7.23 (d, *J* = 4.9 Hz, 1H), 3.72 (s, 2H), 2.16 (s, 3H).

**MS (ESI+)** m/z calculated for C_14_H_14_ClN_2_O^+^ ([M+H]^+^) 261.1, found: 261.0.

**(R)-6-Chloro-N-(isoquinolin-4-yl)chromane-4-carboxamide (Z4643752419) and (S)-6-Chloro-N-(isoquinolin-4-yl)chromane-4-carboxamide (Z4646694589)**

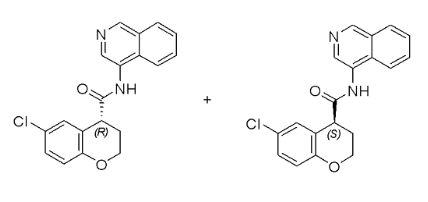

6-chlorochromane-4-carboxylic acid (0.156 g, 735.84 μmol, 1 eq), isoquinolin-4-amine (0.127 g, 937.72 μmol, 1.2 eq), HATU (377.5 mg, 0.99 mmol, 1.35 eq), and DIPEA (284.77 mg, 2.20 mmol, 3 eq) were mixed in anhydrous DMSO (3 mL), and the solution was stirred at 20 °C for 12 h . After that, the reaction mixture was purified by reverse phase HPLC to afford (R,S)-6-chloro-N-(isoquinolin-4-yl)chromane-4-carboxamide (90 mg, 36%).

(R)-6-Chloro-N-(isoquinolin-4-yl)chromane-4-carboxamide (Z4643752419) and (S)-6-Chloro-N-(isoquinolin-4-yl)chromane-4-carboxamide (Z4646694589) were obtained as two fractions (RT = 32.069 min and 46.990 min respectively) from separation of the (R,S)-6-chloro-N-(isoquinolin-4-yl)chromane-4-carboxamide (90 mg) using a Chiralpak IC-III (250*20 mm, 5 μm) column, eluting with hexane:IPA:MeOH (80-10-10) with flow rate of 13 mL/min.

**^1^H NMR (500 MHz, DMSO-*d_6_*)** δ (ppm) 10.38 (s, 1H), 9.17 (s, 1H), 8.69 (s, 1H), 8.16 (d, *J* = 8.2 Hz, 1H), 8.09 (d, *J* = 8.5 Hz, 1H), 7.85 (dd, *J* = 8.4, 7.0 Hz, 1H), 7.72 (t, *J* = 7.5, 7.5 Hz, 1H), 7.37 (d, *J* = 2.6 Hz, 1H), 7.19 (dd, *J* = 8.8, 2.7 Hz, 1H), 6.85 (d, *J* = 8.8 Hz, 1H), 4.43-4.35 (m, 1H), 4.26-4.18 (m, 1H), 4.17 (t, *J* = 5.7, 5.7 Hz, 1H), 2.30-2.20 (m, 2H).

**MS (ESI+)** m/z calculated for C_19_H_16_ClN_2_O_2_^+^ ([M+H]^+^) 339.1, found: 339.0.

**(R,S)-6-chloro-N-(isoquinolin-4-yl)-1,2,3,4-tetrahydroisoquinoline-4-carboxamide**

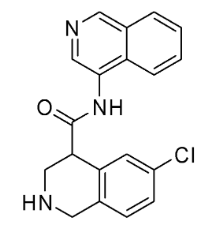

**Step1.** 4-bromo-6-chloroisoquinoline (6.9 g, 28.64 mmol), triethylamine (2.88 g, 28.47 mmol) and Pd(dppf)Cl_2_.2CH_2_Cl_2_ (1.16 g, 1.43 mmol) were dissolved in MeOH (100 mL). The reaction mixture was stirred at 80 °C under CO atmosphere (25 bar) for 16 h. After cooling, the reaction mixture was filtered, and the combined MeOH solution was evaporated under reduced pressure to give a residue. The residue was then suspended in water, and the suspension was filtered, air-dried to afford methyl 6-chloroisoquinoline-4-carboxylate (5.0 g, 79.3%). which was used in the subsequent step without further purification.

**Step 2.** Methyl 6-chloroisoquinoline-4-carboxylate (5.0 g, 22.62 mmol) was dissolved in acetic acid (50 mL). Sodium cyanoborohydride (4.25 g, 67.48 mmol) was then added portionwise to the solution at room temperature, and the reaction mixture was stirred overnight. After that, the reaction mixture was evaporated, and the residue was diluted with water then extracted with CHCl_3_ (2 x 300 mL). The combined organic layer was dried over sodium sulfate and evaporated *in vacuo* to give methyl 6-chloro-1,2,3,4-tetrahydroisoquinoline-4-carboxylate (3.4 g, 66.8%) which was used in subsequent step without further purification.

**Step 3.** di-*tert*-butyl dicarbonate (3.14 g, 14.4 mmol, 1.2 equiv.) was added to the solution of methyl 6-chloro-1,2,3,4-tetrahydroisoquinoline-4-carboxylate (3.4 g, 12.0 mmol) and NaHCO_3_ (5 equiv., aqueous solution) in MeOH (100 mL). The reaction mixture was stirred at 50 °C overnight. Then, the mixture was diluted with water and extracted with chloroform (2 x 50 mL). The combined organic layer was dried over sodium sulfate and evaporated to give 2-tert-butyl 4-methyl 6-chloro-3,4-dihydroisoquinoline-2,4(1H)-dicarboxylate ( 3.4g), which was used in the subsequent step without further purification.

**Step 4.** 2-tert-butyl 4-methyl 6-chloro-3,4-dihydroisoquinoline-2,4(1H)-dicarboxylate (3.4 g, 10.4 mmol) was mixed in MeOH (30 mL) and H_2_O (60 mL), and then NaOH (1.24 g, 31 mmol) was added to the solution. The reaction mixture was heated at 50 °C overnight. Then, the mixture was diluted with water, acidified with NaHSO_4_ to pH 4 and extracted with chloroform (2 x 50 mL). The combined organic layer was dried over sodium sulfate and evaporated to give 2-(tert-butoxycarbonyl)-6-chloro-1,2,3,4-tetrahydroisoquinoline-4-carboxylic acid (2.9 g), which was used in the subsequent step without further purification.

**Step 5.** 2-(tert-butoxycarbonyl)-6-chloro-1,2,3,4-tetrahydroisoquinoline-4-carboxylic acid (4.0 g, 12.86 mmol), isoquinolin-4-amine hydrochloride (2.21 g, 12.26 mmol), (3-[(ethylimino)methylidene]aminopropyl)dimethylamine hydrochloride (3.51 g, 18.39 mmol), N,N-dimethylpyridin-4-amine (298.58 mg, 2.45 mmol), and triethylamine (1.85 g, 18.34 mmol, 2.55 mL) were suspended in DMF, and the reaction mixture was heated at 50 °C overnight. After cooling to room temperature, the mixture was evaporated to give a residue, which was then suspended in water and filtered to give tert-butyl 6-chloro-4-(isoquinolin-4-ylcarbamoyl)-3,4-dihydroisoquinoline-2(1H)-carboxylate (1.45 g, 35.1%), which was used in the subsequent step without further purification.

**Step 6.** tert-butyl 6-chloro-4-(isoquinolin-4-ylcarbamoyl)-3,4-dihydroisoquinoline-2(1H)-carboxylate (1.39 g, 3.18 mmol) was dissolved in MeOH, and then acetyl chloride (748.0 mg, 9.59 mmol, 680.0 µL) was added at to the solution at room temperature. The reaction mixture was heated at 50 °C overnight. Then, the mixture was diluted with water, basified with potassium carbonate to pH 12, and extracted with CHCl_3_. The organic layer was dried over sodium sulfate. The volatiles were removed under reduced pressure to give  (R,S)-6-chloro-N-(isoquinolin-4-yl)-1,2,3,4-tetrahydroisoquinoline-4-carboxamide (0.9 g, 84%).

**^1^H NMR (500 MHz, DMSO-*d_6_*)** δ (ppm) 11.25 (s, 1H), 9.09 (s, 1H), 8.92 (s, 1H), 8.14 (d, *J* = 8.2 Hz, 1H), 8.00 (d, *J* = 8.5 Hz, 1H), 7.86-7.79 (m, 1H), 7.71 (t, *J* = 7.5, 7.5 Hz, 1H), 7.37 (d, *J* = 2.2 Hz, 1H), 7.26 (dd, *J* = 8.2, 2.3 Hz, 1H), 7.18 (d, *J* = 8.3 Hz, 1H), 4.04 (d, *J* = 16.3 Hz, 1H), 3.92 (d, *J* = 16.4 Hz, 1H), 3.83 (t, J = 4.0, 4.0 Hz, 1H), 3.46 (dd, J = 12.7, 3.5 Hz, 1H), 3.13 (dd, *J* = 12.8, 4.4 Hz, 1H).

**MS (ESI+)** m/z calculated for C_19_H_17_ClN_3_O^+^( [M+H]^+^) 338.1,  found: 338.2.

**rel-(R)-6-chloro-N-(isoquinolin-4-yl)-1,2,3,4-tetrahydroisoquinoline-4-carboxamide (Z4943052515, rel-BEN-DND-f2e727cd-5-1) and rel-(R)-6-chloro-N-(isoquinolin-4-yl)-1,2,3,4-tetrahydroisoquinoline-4-carboxamide (Z4943052518, rel-BEN-DND-f2e727cd-5-2)**

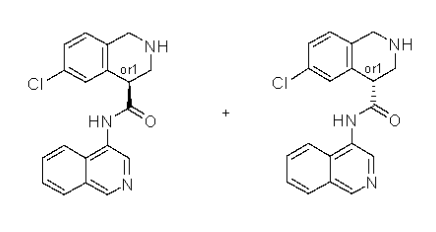

(R,S)-6-chloro-N-(isoquinolin-4-yl)-1,2,3,4-tetrahydroisoquinoline-4-carboxamide (50 mg) was separated using the condition mentioned below to afford two isomers (**Z4943052515, Z4943052518**) at RT = 21.465 min and RT = 31.200 min.

Chiral separation condition:

Column: Chiralcel OD-H (250*20 mm, 5 µm)

Mobile phase: hexane:IPA:MeOH, 80:10:10

Flow rate: 12 mL/min

***Z4943052515, rel-BEN-DND-f2e727cd-5-1:***

Yield: 18.02 mg (36.04%)

Analytical RT = 11.878 min (Chiralcel OD-H (250*4.6 mm, 5 µm), hexane:IPA:MeOH, 70:15:15, 0.6 mL/min) .

***Z4943052518, rel-BEN-DND-f2e727cd-5-2:***

Yield: 17.94 mg (35.88%)

Analytical RT = 15.304 min (Chiralcel OD-H (250*4.6 mm, 5 µm), hexane:IPA:MeOH, 70:15:15, 0.6 mL/min).

**(R,S)-6-Chloro-N-(isoquinolin-4-yl)-2-(methylsulfonyl)-1,2,3,4-tetrahydroisoquinoline-4-carboxamide**

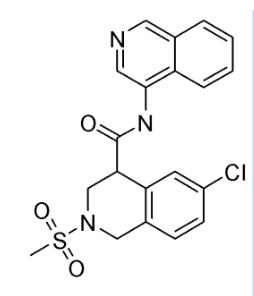

6-chloro-N-(isoquinolin-4-yl)-1,2,3,4-tetrahydroisoquinoline-4-carboxamide (100 mg, 0.3 mmol) and DIPEA (0.1 mL) were dissolved in DMF (4 mL), and then methanesulfonyl chloride (40 mg, 0.36 mmol) was added to the solution. The reaction mixture was stirred at room temperature for 16 h. Then, the mixture was purified by HPLC (25-50% 0-5 min H_2_O/ACN, flow: 30 mL/min (loading pump 4 mL/min acetonitrile); column: Chromatorex 18 SMB100-5T 100x19 mm 5 µm) to give (R,S)-6-chloro-N-(isoquinolin-4-yl)-2-(methylsulfonyl)-1,2,3,4-tetrahydroisoquinoline-4-carboxamide (34 mg).

**^1^H NMR (500 MHz, DMSO-*d_6_*)** δ (ppm) 10.37 (s, 1H), 9.18 (s, 1H), 8.67 (s, 1H), 8.16 (dd, *J* = 8.3, 4.9 Hz, 2H), 7.87-7.80 (m, 1H), 7.72 (t, *J* = 7.5, 7.5 Hz, 1H), 7.42 (d, *J* = 2.1 Hz, 1H), 7.35 (dd, *J* = 8.3, 2.2 Hz, 1H), 7.31 (d, *J* = 8.3 Hz, 1H), 4.43 (s, 2H), 4.36 (t, *J* = 5.9, 5.9 Hz, 1H), 3.83 (d, J = 5.8 Hz, 2H), 3.00 (s, 3H).

MS (ESI+) m/z calculated for C_20_H_19_ClN_3_O_3_S^+^([M+H]^+^) 416.1, found: 416.0.

**rel-(R)-6-chloro-N-(isoquinolin-4-yl)-2-(methylsulfonyl)-1,2,3,4-tetrahydroisoquinoline-4-carboxamide (Z4988872945) and rel-(R)-6-chloro-N-(isoquinolin-4-yl)-2-(methylsulfonyl)-1,2,3,4-tetrahydroisoquinoline-4-carboxamide (Z4988873021)**

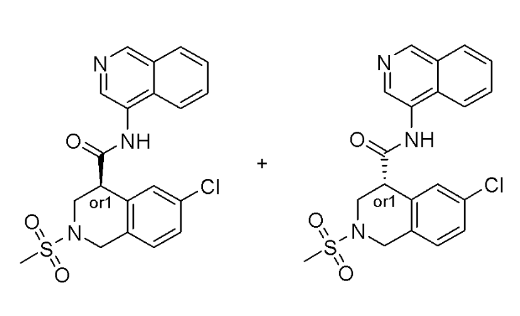

(R,S)-6-Chloro-N-(isoquinolin-4-yl)-2-(methylsulfonyl)-1,2,3,4-tetrahydroisoquinoline-4-carboxamide (50 mg) was separated using the condition mentioned below to afford two isomers (***Z4988872945 and Z4988873021***) at RT = 41.218 min and RT = 51.863 min.

**Chiral separation condition:**

Column: Chiralpak IG (250*20 mm, 5 µm)

Mobile phase: hexane:IPA:MeOH, 50:25:25

Flow rate: 12 mL/min

***Z4988872945:***

Yield: 10.49 mg (42.30%)

Analytical RT = 20.869 min (Chiralpak IG (250*4.6 mm, 5 µm), IPA:MeOH, 50:50, 0.6 mL/min).

***Z4988873021:***

Yield: 10.40 mg (41.94%)

Analytical RT = 16.342 min (Chiralpak IG (250*4.6 mm, 5 µm), IPA:MeOH, 50:50, 0.6 mL/min).

**(R,S)-6-chloro-2-(((1-cyanocyclopropyl)methyl)sulfonyl)-N-(isoquinolin-4-yl)-1,2,3,4-tetrahydroisoquinoline-4-carboxamide**

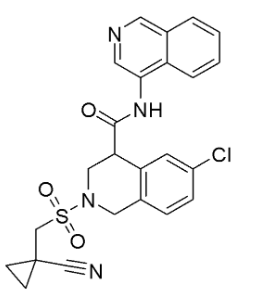

**Step 1.** tert-butyl 6-chloro-4-(isoquinolin-4-ylcarbamoyl)-3,4-dihydroisoquinoline-2(1H)-carboxylate (2.2 g, 97% purity) was dissolved in dioxane, and HCl (8 % solution in dioxane, 3 equiv.) was added to the solution. The reaction mixture was stirred at room temperature overnight. The crystalline substance was filtered off, washed with acetone, and dried to afford crude 6-chloro-N-(isoquinolin-4-yl)-1,2,3,4-tetrahydroisoquinoline-4-carboxamide dihydrochloride, which was used in the next step without further purification (1.8 g, 98%).

**Step 2.** 6-chloro-N-(isoquinolin-4-yl)-1,2,3,4-tetrahydroisoquinoline-4-carboxamide dihydrochloride (600.0 mg, 1.47 mmol) and triethylamine (739.52 mg, 7.31 mmol, 1.02 mL) were dissolved in DCM and cooled to 5-10 °C. (1-cyanocyclopropyl)methanesulfonyl chloride (393.82 mg, 2.2 mmol) was then added to the reaction, and the reaction mixture was stirred at room temperature for 48 h. The reaction mixture was evaporated, and the crude product was purified by preparative HPLC to afford (R,S)-6-chloro-2-(((1-cyanocyclopropyl)methyl)sulfonyl)-N-(isoquinolin-4-yl)-1,2,3,4-tetrahydroisoquinoline-4-carboxamide (142.0 mg, 20.2%).

**^1^H NMR (500 MHz, DMSO-*d_6_*)** δ (ppm) 10.43 (s, 1H), 9.18 (s, 1H), 8.68 (s, 1H), 8.16 (t, *J* = 8.5, 8.5 Hz, 2H), 7.84 (t, *J* = 7.7, 7.7 Hz, 1H), 7.72 (t, *J* = 7.5, 7.5 Hz, 1H), 7.43 (d, *J* = 2.1 Hz, 1H), 7.39-7.33 (m, 1H), 7.31 (d, *J* = 8.4 Hz, 1H), 4.57-4.46 (m, 2H), 4.36 (t, *J* = 6.0, 6.0 Hz, 1H), 3.97-3.84 (m, 2H), 3.61-3.50 (m, 2H), 1.43-1.38 (m, 2H), 1.24-1.19 (m, 2H).

**MS (ESI+)** m/z calculated for C_24_H_22_ClN_4_O_3_S^+^([M+H]^+^) 481.1, found: 481.2.

**(4S)-6-chloro-2-[(1-cyanocyclopropyl)methylsulfonyl]-N-(4-isoquinolyl)-3,4-dihydro-1H-isoquinoline-4-carboxamide (Z5129808241, rel-MAT-POS-dc2604c4-1-2) and (4R)-6-chloro-2-[(1-cyanocyclopropyl)methylsulfonyl]-N-(4-isoquinolyl)-3,4-dihydro-1H-isoquinoline-4-carboxamide (Z5129808244, re-MAT-POS-dc2604c4-1-1)**

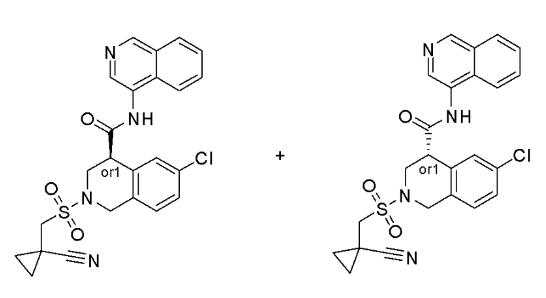

(R,S)-6-chloro-2-(((1-cyanocyclopropyl)methyl)sulfonyl)-N-(isoquinolin-4-yl)-1,2,3,4-tetrahydroisoquinoline-4-carboxamide (90 mg) was separated using the condition mentioned below to afford (***Z4988872945 and Z4988873021***) at RT = 15.399 min and 23.429 min.

**Chiral separation condition:**

Column: CHIRALPAK IС (250x21 mm, 5 µm)

Mobile phase: IPA:MeOH, 50:50

Flow rate: 12 mL/min

***Z5129808241, rel-MAT-POS-dc2604c4-1-2:***

Yield: 480.37 mg (48.04%)

Analytical RT = 12.281 min (Chiralpak IA (250x4.6 mm, 5 µm); MeOH:IPA, 50:50; flow rate: 0.6 mL/min).

***Z5129808244, rel-MAT-POS-dc2604c4-1-1:***

Yield: 436.07 mg (43.61%)

Analytical RT = 20.221 min (Chiralpak IA (250x4.6 mm, 5 µm); MeOH:IPA, 50:50; flow rate: 0.6 mL/min).

**(2-hydroxyquinolin-4-yl)(4-(3-(trifluoromethyl)phenyl)piperazin-1-yl)methanone**

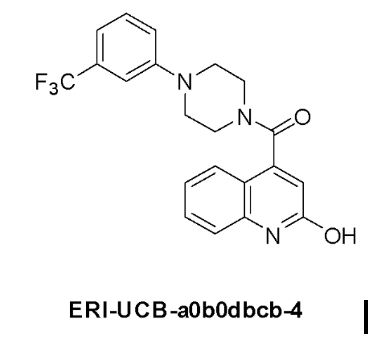

A vial was charged with 2-hydroxyquinoline-4-carboxylic acid (typically 0.1 mmol, 1.0 equiv.), and CDI (prepared in advance at 15% solution in DMSO, 1.1 equiv.) was then added to the vial. The reaction mixture was heated with stirring at 50 °C for 2 h. After that, 1-(3-(trifluoromethyl)phenyl)piperazine (1.1 equiv.) was added to the reaction, and the vial was sealed and heated at 100 °C for 6 h. In the case of using a salt of the amine, an additional amount of DIPEA was added to the reaction mixture to convert the amine to the basic form. After cooling to the ambient temperature, the mixture was filtered and the solution was subjected to HPLC purification.

**^1^H NMR (600 MHz, DMSO-*d_6_*)** δ (ppm) 11.92 (s, 1H), 7.53 (m, 1H), 7.42 (m, 1H), 7.34 (m, 1H), 7.19 (m, 3H), 7.07 (d, *J* = 5.0 Hz, 1H), 6.51 (s, 1H), 3.80 (m, 2H), 3.32 (m, 4H), 3.20 (m, 1H), 3.01 (m, 1H).

**MS (ESI+)** m/z calculated for C_21_H_19_F_3_N_3_O_2_^+^ ([M+H]^+^) 402.1, found: 402.4.

**Separation condition:**

Column: Waters Sunfire C18 OBD Prep Column, 100 Å^2^, 5 µm, 100x19 mm with SunFire C18 Prep Guard Cartridge

Mobile phase: H_2_O:MeOH. In some cases, ammonia or TFA was used as an additive to improve the separation of the products.

**6-chloro-*N*-(isoquinolin-4-yl)thiochromane-4-carboxamide 1,1-dioxide**

**Step 1.** To the solution of 6-chloro-3,4-dihydro-2H-1-benzothiopyran-4-one (2.0 g, 10.07 mmol)  in DCM (20 mL) were added trimethylsilanecarbonitrile (1.3 g, 13.09 mmol)  and a catalytic amount of diiodozinc (160.69 mg, 503.4 µmol). The reaction mixture was stirred at room temperature overnight, at which time, a check by NMR showed full conversion. The volatiles were removed in vacuo to give a crude 6-chloro-4-[(trimethylsilyl)oxy]-3,4-dihydro-2H-1-benzothiopyran-4-carbonitrile (2.99 g, 99.7%), which was used in the next step without further purification.

**Step 2.** To a 50-mL single neck round bottom reaction flask equipped with a magnetic stirrer, a reflux condenser, and a nitrogen inlet were added 6-chloro-4-[(trimethylsilyl)oxy]-3,4-dihydro-2H-1-benzothiopyran-4-carbonitrile (2.99 g, 10.04 mmol), tin(II) chloride dihydrate (9.14 g, 40.16 mmol), glacial acetic acid (8 mL), and concentrated hydrochloric acid ( 8 mL). The reaction apparatus was immediately flushed with Argon and plunged into a preheated (100 ^o^C) oil bath. With vigorous stirring, the reaction mixture was heated for 48 h. After cooling to room temperature, the mixture was diluted with water (20 mL) and extracted with DCM (2 x 20 mL). The combined organic layer was dried over Na_2_SO_4_ and evaporated to give a residue, which was then suspended in NaOH (1 N, 30 mL)  and DCM (20 mL). The aqueous layer was carefully acidified with HCl (10%) to pH = 2 and then extracted with DCM (3 x 15 mL). The organic layers were combined, dried over Na_2_SO_4_, and then evaporated to give 6-chloro-3,4-dihydro-2H-1-benzothiopyran-4-carboxylic acid (1.44 g, 62.7%)

**Step 3.** To a mixture of 6-chloro-3,4-dihydro-2H-1-benzothiopyran-4-carboxylic acid (1.0 g, 4.37 mmol) in acetonitrile:H_2_O (1:1, 20 mL) was added trichlororuthenium hydrate (19.72 mg, 87.46 µmol). After that, sodium periodate (2.34 g, 10.93 mmol) was added in portions. The reaction mixture was then stirred for 3 h. Solids were filtered and washed with water (10 mL) and ethyl acetate (25 ml). The organic layer was separated, and the aqueous layer was re-extracted with ethyl acetate (2 x 10 mL). The combined organic layer was dried over Na_2_SO_4_ and evaporated to give a residue, which was then dissolved in acetonitrile (25 mL), and SiliaMetS Thiol Metal Scavengers was added to the solution. The mixture was stirred for 30 min and then filtered. The filtrate was evaporated in vacuo to give 6-chloro-1,1-dioxo-3,4-dihydro-2H-1lambda6-benzothiopyran-4-carboxylic acid (940.0 mg, 82.5%).

**Step 4.** DIPEA (86.76 mg, 671.32 µmol) was added to a mixture of 6-chloro-1,1-dioxo-3,4-dihydro-2H-1lambda6-benzothiopyran-4-carboxylic acid (50.0 mg, 191.8 µmol), isoquinolin-4-amine hydrochloride (41.58 mg, 230.17 µmol), and HATU (109.39 mg, 287.71 µmol) in DMF (2 mL). The reaction mixture was stirred at room temperature overnight. The reaction progress was checked by LCMS. After consumption of starting material, the reaction mixture was subjected to prepHPLC to give 6-chloro-N-(isoquinolin-4-yl)-1,1-dioxo-3,4-dihydro-2H-1lambda6-benzothiopyran-4-carboxamide (42.8 mg, 57.7%).

**MS (ESI+)** m/z calculated for C_19_H_16_ClN_2_O_3_S^+^ ([M+H]^+^) 387.1, found: 387.2.

**Separation condition:**

Column: Chromatorex 18 SMB100-5T 100x19 mm 5 µm,

Mobile phase: 20-35% in 0-6 min, H_2_O/acetonitrile)

Flow rate: 30 mL/min

**6,7-dichloro-N-(isoquinolin-4-yl)-4-methyl-1,2,3,4-tetrahydroquinoline-4-carboxamide**

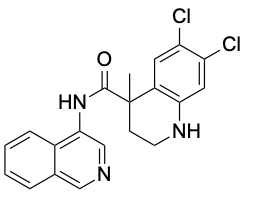

**Step 1.** Methyl 6,7-dichloro-1,2,3,4-tetrahydroquinoline-4-carboxylate (1.98 g, 7.6 mmol) in anhydrous THF (20 mL) was added dropwise at -30 °C under a nitrogen atmosphere to a solution of lithium diisopropylamide (1 equiv.)  in THF. The mixture was stirred at room temperature for 1 h. After that, iodomethane (1.62 g, 11.4 mmol, 710.0 µL, 1.5 equiv.) was added at -50 °C to the mixture, and then the reaction was stirred at room temperature overnight. The reaction mixture was quenched with saturated aqueous NH_4_Cl and extracted with ethyl acetate. The organic layer was washed with saturated brine, dried over anhydrous Na_2_SO_4_, and evaporated in vacuo to give a residue, which was purified by column chromatography (SiO_2_, hexane/ethyl acetate) to give methyl 6,7-dichloro-4-methyl-1,2,3,4-tetrahydroquinoline-4-carboxylate (400.0 mg, 19.2%).

**Step 2.** Methyl 6,7-dichloro-4-methyl-1,2,3,4-tetrahydroquinoline-4-carboxylate (400.0 mg, 1.46 mmol) was dissolved in methanol (15 mL), and then sodium hydroxide (70.12 mg, 1.75 mmol) in water (8 mL) was added to the solution. The obtained mixture was stirred at room temperature overnight, concentrated in vacuo to give sodium 6,7-dichloro-4-methyl-1,2,3,4-tetrahydroquinoline-4-carboxylate (400.0 mg, 97.5%)

**Step 3.** To a suspension of sodium 6,7-dichloro-4-methyl-1,2,3,4-tetrahydroquinoline-4-carboxylate (400.0 mg, 1.42 mmol)  in dioxane (8 mL) and H_2_0 (8 mL) was added sodium hydrogen carbonate (357.32 mg, 4.25 mmol) and prop-2-en-1-yl carbonochloridate (256.34 mg, 2.13 mmol, 230.0 µL, 1.5 equiv.). After stirring overnight at room temperature, the reaction mixture was washed with methyl tertiary-butyl ether. Hydrochloric acid was then added to the water layer,  and the mixture was extracted with ethyl acetate. The organic layer was washed with saturated brine, dried over anhydrous Na_2_SO_4_, and evaporated in vacuo to give 6,7-dichloro-4-methyl-1-[(prop-2-en-1-yloxy)carbonyl]-1,2,3,4-tetrahydroquinoline-4-carboxylic acid (300.0 mg, 61.5%).

**Step 4.** HATU (321.66 mg, 845.97 µmol)  was added to a mixture of  6,7-dichloro-4-methyl-1-[(prop-2-en-1-yloxy)carbonyl]-1,2,3,4-tetrahydroquinoline-4-carboxylic acid (253.2 mg, 735.63 µmol),  isoquinolin-4-amine (106.06 mg, 735.63 µmol), and ethylbis(propan-2-yl)amine (237.44 mg, 1.84 mmol, 320.0 µL, 2.5 equiv.)  in DMF (10 mL). The obtained mixture was stirred at room temperature overnight, then poured into water, and extracted with ethyl acetate.  The organic layer was washed with saturated brine, dried over anhydrous Na_2_SO_4_, and evaporated in vacuo to give a residue, which was purified by HPLC (Chromatorex (18 SMB100-5T 100x19 mm), H_2_O/acetonitrile) to give prop-2-en-1-yl 6,7-dichloro-4-[(isoquinolin-4-yl)carbamoyl]-4-methyl-1,2,3,4-tetrahydroquinoline-1-carboxylate (14.0 mg, 4%) .

**Step 5.** To a solution of prop-2-en-1-yl 6,7-dichloro-4-[(isoquinolin-4-yl)carbamoyl]-4-methyl-1,2,3,4-tetrahydroquinoline-1-carboxylate (14.0 mg, 29.77 µmol)  in DCM (5 mL) under argon were added Pd(PPh_3_)4 (3.45 mg, 2.98 µmol) and morpholine (5.2 mg, 59.69 µmol, 10.0 µL, 2.0 equiv.) . After stirring overnight at room temperature, the obtained mixture was concentrated in vacuo and then subjected to separation (Chiralpak AS-H (250*20 mm, 5 µm), hexane-IPA-MeOH) without additional work-up to give 6,7-dichloro-N-(isoquinolin-4-yl)-4-methyl-1,2,3,4-tetrahydroquinoline-4-carboxamide (9.91 mg, 86.2% yield).

**MS (ESI+)** m/z calculated for C_20_H_18_Cl_2_N_3_O^+^ ([M+H]) 386.1, found: 386.0.

**6-chloro-N-(isoquinolin-4-yl)-3-methyl-3,4-dihydro-2H-1-benzopyran-4-carboxamide**

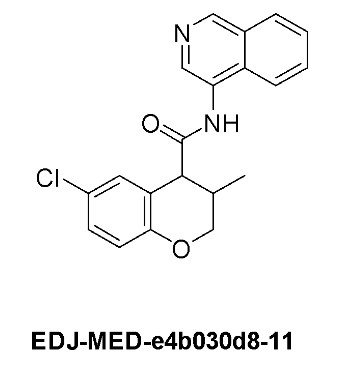

To a solution of 6-chloro-3-methyl-3,4-dihydro-2H-1-benzopyran-4-carboxylic acid (1.1 g, 4.85 mmol) in DMF (5 mL) were added isoquinolin-4-amine hydrochloride (876.51 mg, 4.85 mmol), DIPEA (1.25 g, 9.7 mmol, 1.69 mL, 2.0 equiv.) and HATU (2.03 g, 5.34 mmol), and the reaction mixture was stirred overnight at room temperature. Water (30 mL) was then added to the reaction, and the solution was extracted with ethyl acetate (4 x 50 mL). The combined organic layers were washed with brine, dried with sodium sulfate, filtered, and evaporated in vacuo to give a residue, which was purified by flash chromatography using a CombiFlash to afford 6-chloro-N-(isoquinolin-4-yl)-3-methyl-3,4-dihydro-2H-1-benzopyran-4-carboxamide (1.2 g, 93.0% purity, 65.2%).

**^1^H NMR (600 MHz, DMSO-*d_6_*)** δ (ppm) 10.42 (s, 1H), 9.12 (s, 1H), 8.66 (s, 1H), 8.18-8.11 (m, 2H), 7.90-7.88 (m, 1H), 7.73-7.71 (m, 1H), 7.32 (s, 1H), 7.19-7.17 (m, 1H), 6.83 (d, *J* = 6.0 Hz, 1H), 4.31-4.28 (m, 1H), 4.10 - 4.07 (m, 2H), 2.32-2.30 (m, 1H), 1.07 (d, *J* = 8.4 Hz, 3H).

**MS (ESI+)** m/z calculated for C_20_H_18_ClN_2_O_2_^+^ ([M+H]^+^) 353.1, found: 353.0.

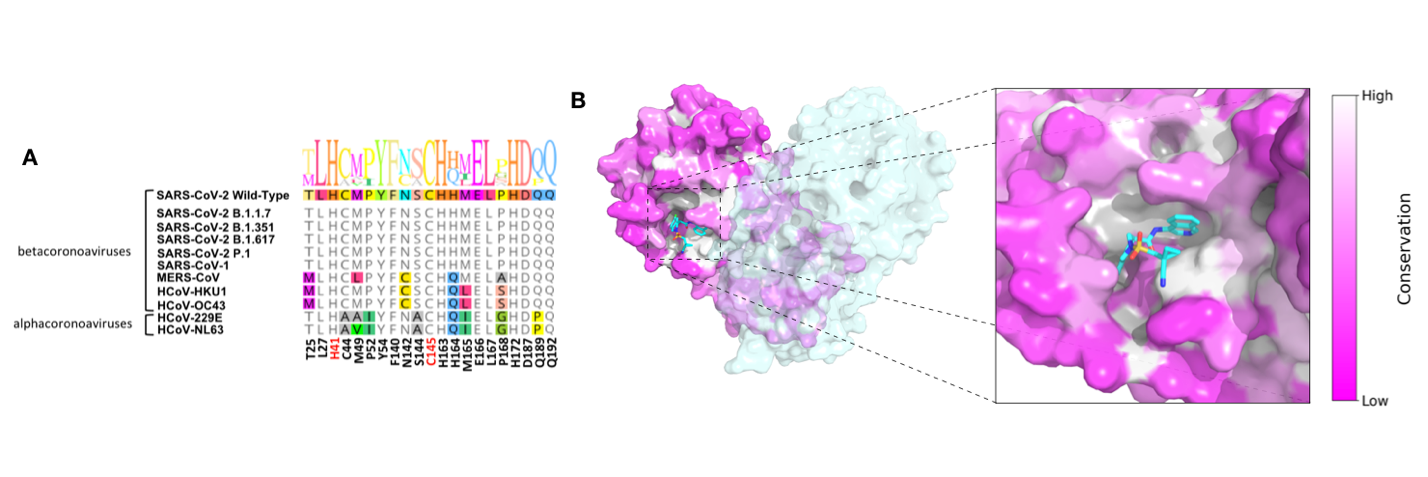

Fig. S1. The SARS-CoV-2 main viral protease (Mpro) is highly conserved across coronaviruses. A. Mpro sequences across coronaviruses are highly conserved due to their requirement to cleave viral polyproteins in numerous locations, showing very little variation in residues lining the active site near the scissile bond. B. Available structural data for Mpro from multiple coronaviruses shows a high degree of sequence and structural conservation, especially in the vicinity of the active site.

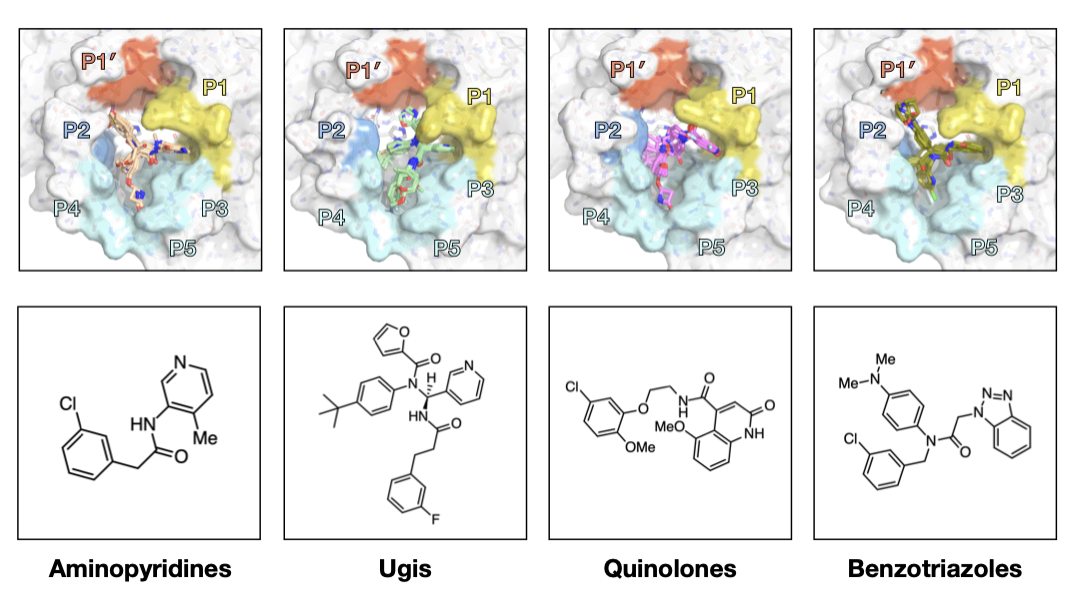

Fig. S2. Initial crowdsourcing efforts produced structures and hits with biochemical potency for four distinct chemical series. Ugi, quinolones and benzotriazoles were based on literature compounds for SARS-CoV main protease. *Top:* Early representative X-ray structures of each series. *Bottom*: Representative compounds from each series.

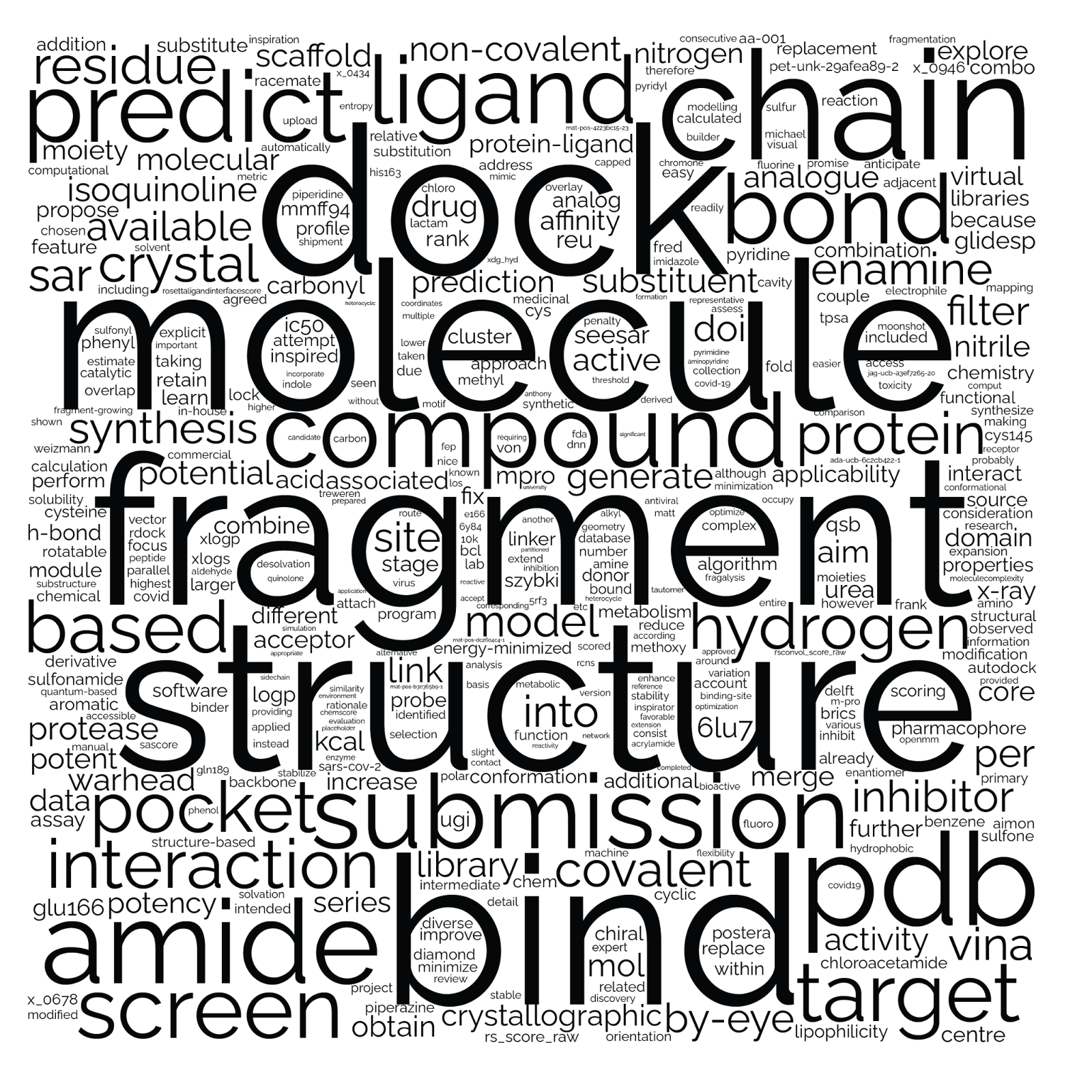

**Fig S3. Word cloud representation of crowdsourced submission description.** Representation of word frequency from 2,127 crowdsourced submission descriptions. Although this is just a qualitative assessment, some trends may emerge (beyond the obvious theme of **structure** based **fragment** optimization) for instance many of the predictions relied on docking (“**dock**”, “**vina**”) and less were “**by-eye**”) several of the predictions attempted “**link**” “**combine**” or “**merge**” presumably fragments.

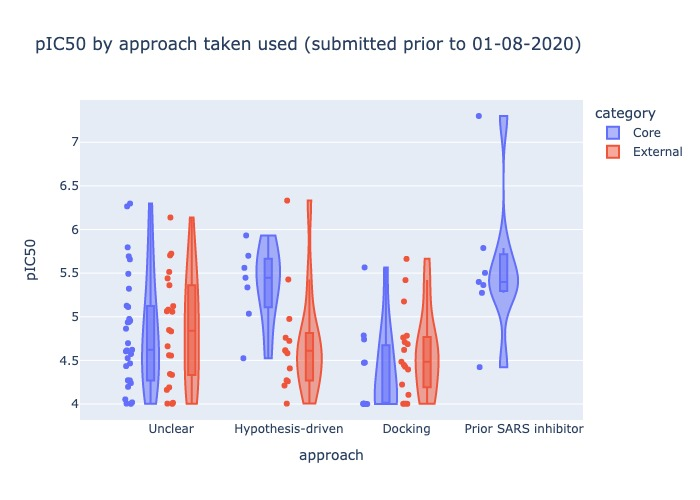

**Fig S4. Breakdown of early submitted designs by methods.** We attempted to evaluate if a specific ‘design methods’ was more successful based on the descriptions of early designs. It is clear the historical SARS inhibitors perform the best (therefore it might be best to start from a known inhibitor when one is available). Other approaches however do not give a clear signal. While Hypothesis-driven designs seem to be more potent than ‘docking’ driven designs, this is driven mainly by a group of covalent designs submitted by the core group.

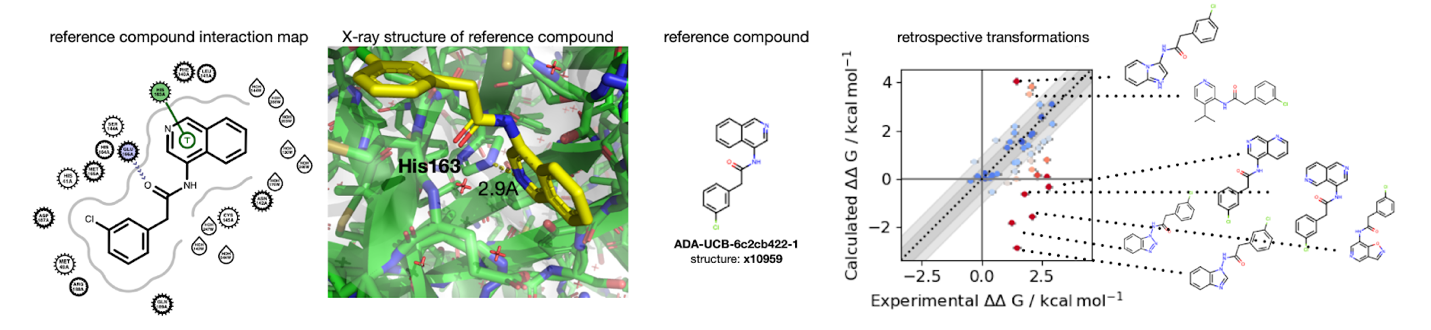

**Fig S5. Outliers in retrospective alchemical free energy calculations drove new chemical insight that informed improvements in modeling strategy.** Alchemical free energy calculation [Sprint 10](https://fah-public-data-covid19-moonshot-sprints.s3.us-east-2.amazonaws.com/dashboards/sprint-10/sprint-10-2021-07-26-x10959-dimer-neutral-restrained/retrospective_microstate_transformations/index.html) considered transformations proposed from **ADA-UCB-6c2cb422-1** as a reference compound. [Retrospective calculations](https://fah-public-data-covid19-moonshot-sprints.s3.us-east-2.amazonaws.com/dashboards/sprint-10/sprint-10-2021-07-26-x10959-dimer-neutral-restrained/retrospective_microstate_transformations/index.html) included in the batch showed excellent correspondence for many compounds but a near-vertical band of outliers with no correlation with experiment. Examination of the outliers in this band (right) revealed they shared conservative modifications to the quinoline scaffold engaging P1 that made very small changes to steric contacts but significantly modified the pKa of the nitrogen engaging His163, suggesting we needed to consider multiple protonation state variants of both His163 and the part of the compound engaging it in hydrogen bonding. Future iterations of free energy calculations considered multiple protonation states of His163 and design compounds.

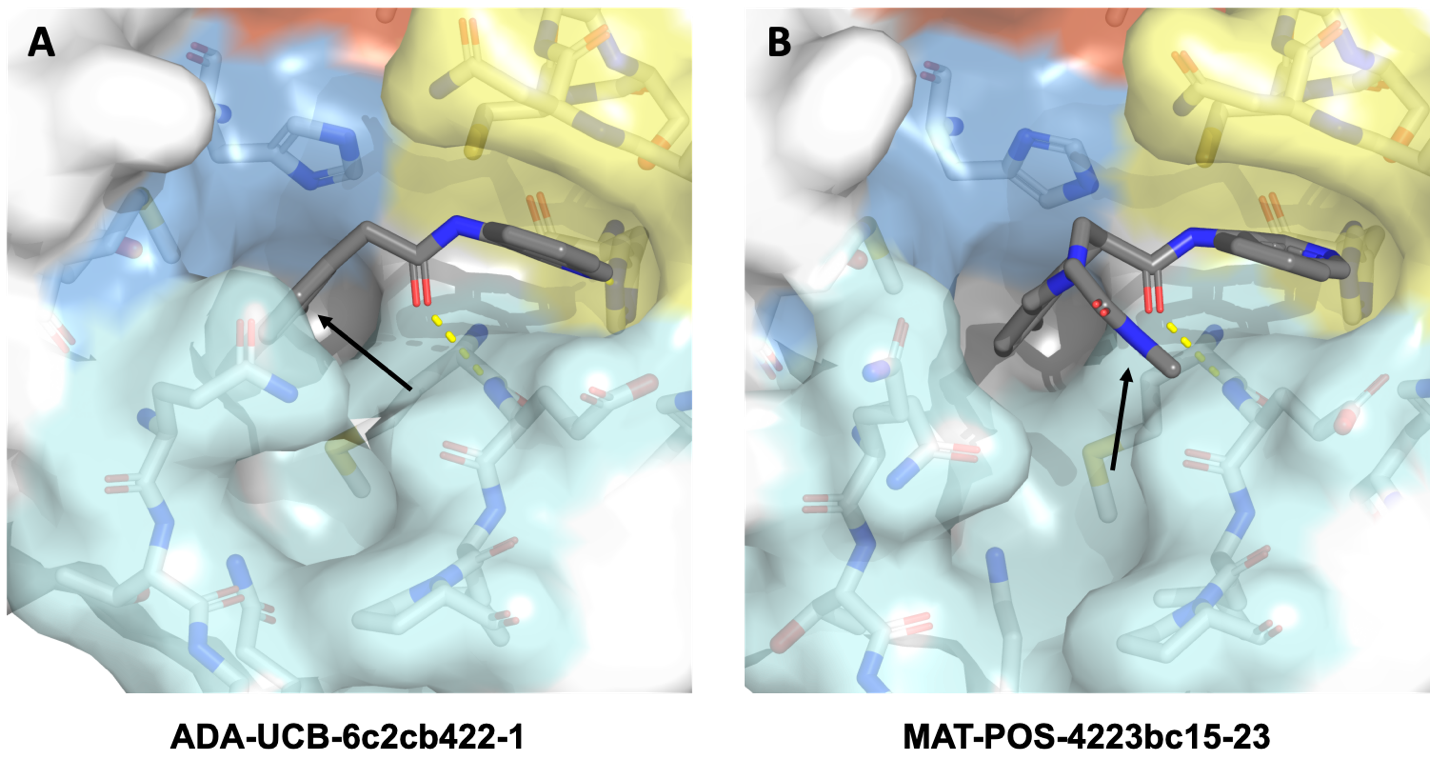
 **Fig S6. Parent compounds for HTC optimization. A.** Structure of the precursor for the Chan-Lam reaction, in which we attempted to functionalize Position C5 of the chloro-phenyl (marked with an arrow) to expand into the P3-P5 pockets (cyan). PDB ID: 7GLP; Resolution 1.92Å **B.** Structure of example of a methyl-derived amide for the parent acid **MAT-POS-4223bc15-21** (for which we were not able to get a good co-crystal structure, possibly due to its relatively weak affinity). We attempted to derivatize the amide bond to expand towards the P3-P5 pockets (cyan). PDB ID: 7GKV; Resolution 1.88Å.

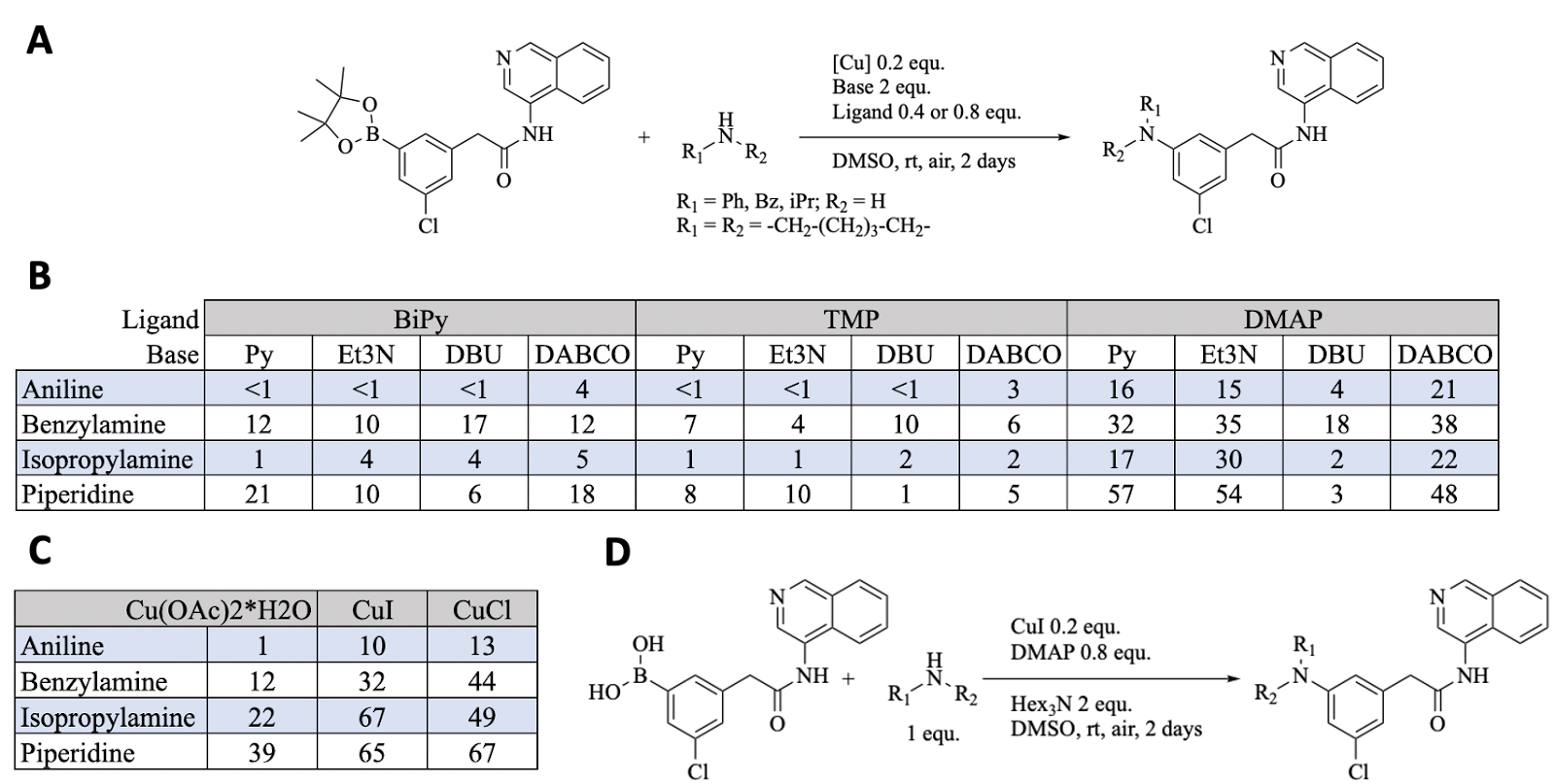

**Fig S7. Optimization of HTC Chan-Lam Reaction. A.** Model reaction scheme. **B.** Initial reaction condition screening was performed to find the most suitable base (Py: Pyridine; Et_3_N: triethylamine; DBU: 1,8-Diazabicyclo(5.4.0)undec-7-ene; DABCO: 1,4-diazabicyclo[2.2.2]octane) and ligand (BiPy: Bipyridine; TMP: 3,4,7,8-Tetramethyl1,10-phenanthroline; DMAP: 4-Dimethylaminopyridine) for the synthesis of arylamines using DMSO as a solvent, atmospheric oxygen as an oxidant, and Cu(OAc)2*H_2_O as a copper source. The experiment was done in a 384-well plate and the reactants were arrayed manually. We report estimated yields (%). Under the tested conditions, the yields correlate with the nucleophilicity of the tested model amines. The highest yields were obtained using DMAP and Et_3_N. **C.** We investigated the effect of copper source on product yield, using the previously optimized reaction conditions (DMAP, Et_3_N). There was no significant difference between copper-iodide (CuI) and copper-chloride (CuCl), thus we decided to proceed with CuI since it is a known promoter for this reaction in combination with DMAP(*93*). **D.** The final optimization step included switching the pinacol boronic ester to an unprotected boronic acid as well as the Et_3_N to trihexylamine - a less volatile homolog of Et_3_N, which yielded 76% product formation when piperidine was used as the model amine, without phenol or homocoupling side-products and 4% of protodeboronated starting material. These reaction conditions were used for library synthesis.

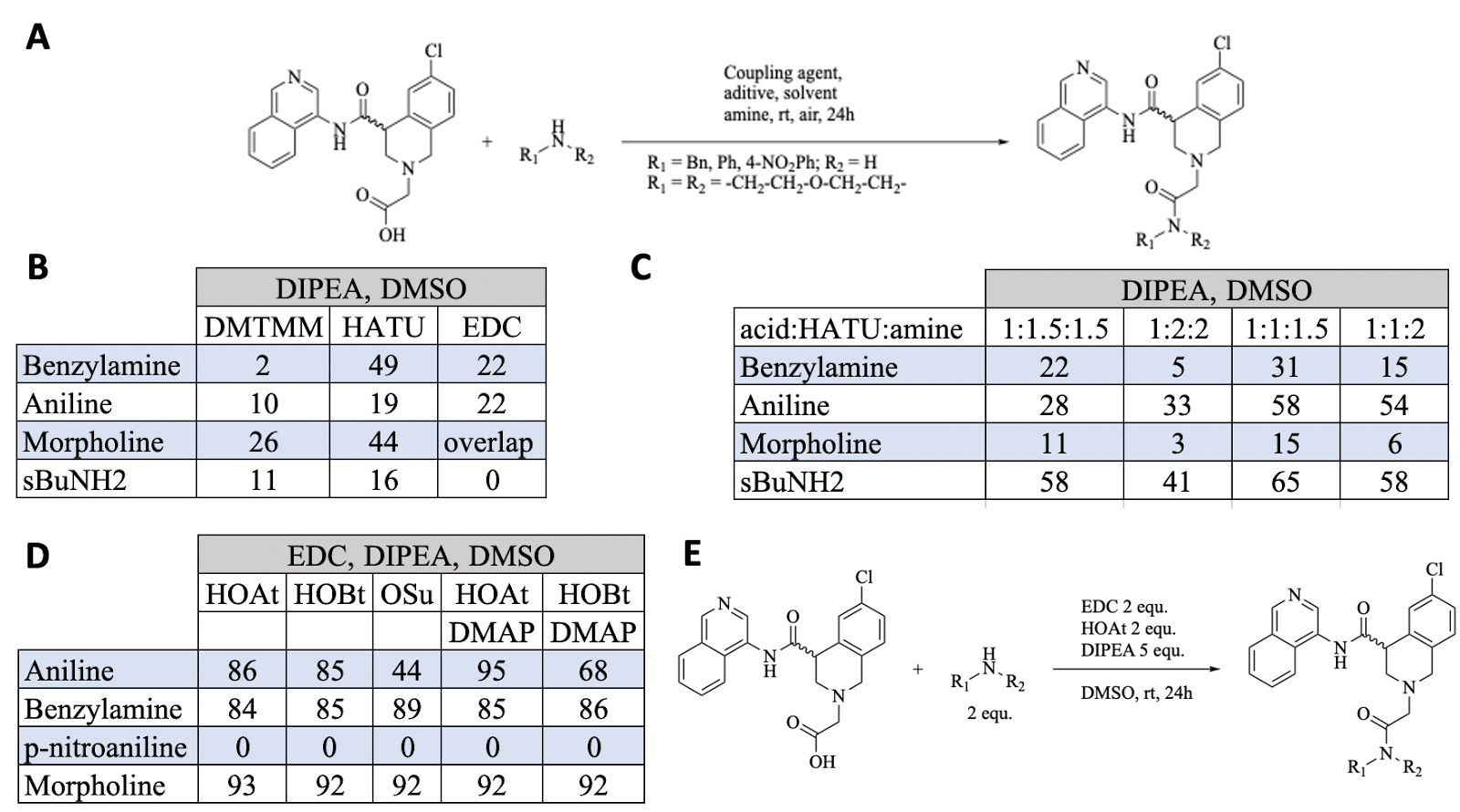

**Fig S8. Optimization of HTC amide coupling. A.** Model reaction scheme. **B.** We investigated suitable coupling agents. Reactions in DMSO, with equimolar amounts of DIPEA (N,N-Diisopropylethylamine), coupling agent (DMT-MM: 4-(4,6-dimethoxy-1,3,5-triazin-2-yl)-4-methyl- morpholinium chloride; EDC: 1-Ethyl-3- (3-dimethylaminopropyl) carbodiimide HCl; HATU: 1-[Bis(dimethylamino) methylene]-1H-1,2,3-triazolo[4,5-b] pyridinium 3-oxide hexafluorophosphate), model amine (aniline, benzylamine, morpholine, and sec-butylamine), and carboxylic acid each. After pre-activating the acid with the coupling agent for five minutes, model amines were arrayed to each well. After overnight incubation, each well was diluted with 50% ACN (Acetonitrile) in water, and the yields were estimated from the UV chromatograms. We report estimated yields (%). In all cases, unreacted acid was observed. **C.** With higher ratios of coupling agent and base, the yields did increase, but uronium modified starting material was observed**.** We tried repeating the reaction in DMA (Dimethylacetamide) or NMP (N-methylpyrrolidone), but the yields were even lower (data not shown). **D.** We increased the amount of DIPEA to 5 equ. and replaced HATU with EDC to avoid formation of side products. Using EDC, we observed 100% acid consumption. To avoid isomerization of the activated acid to unreactive N-acylurea, we explored the use of additives (HOAt: 1-Hydroxy-7-azabenzotriazole; HOBt: Hydroxybenzotriazole; OSu:N,N-Disuccinimidyl carbonate). We report estimated yields (%). **E.** Since DMAP did not significantly improve the yield, we decided to proceed with HOAt as an additive to simplify the UV chromatograms. These reaction conditions were used for library synthesis.

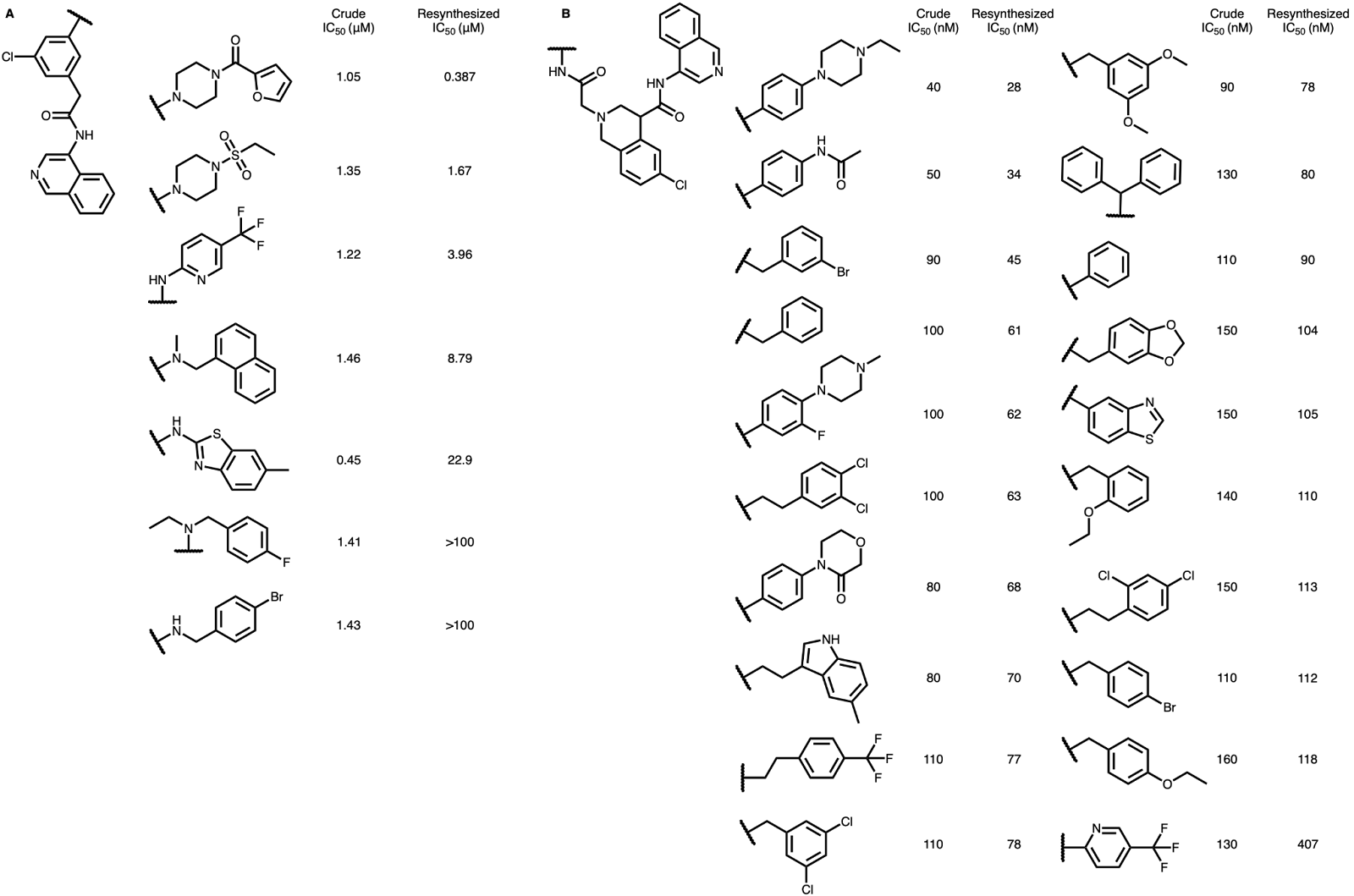

Fig S9. HTC optimization of Mpro inhibitors towards P3-5. A. Selected Chan-Lam installed derivatives of ADA-UCB-6c2cb422-1 and B. Amide coupling derivatives of MAT-POS-4223bc15-23 were tested both as crude reaction mixtures, as well as pure resynthesized compounds in the Mpro biochemical assay.

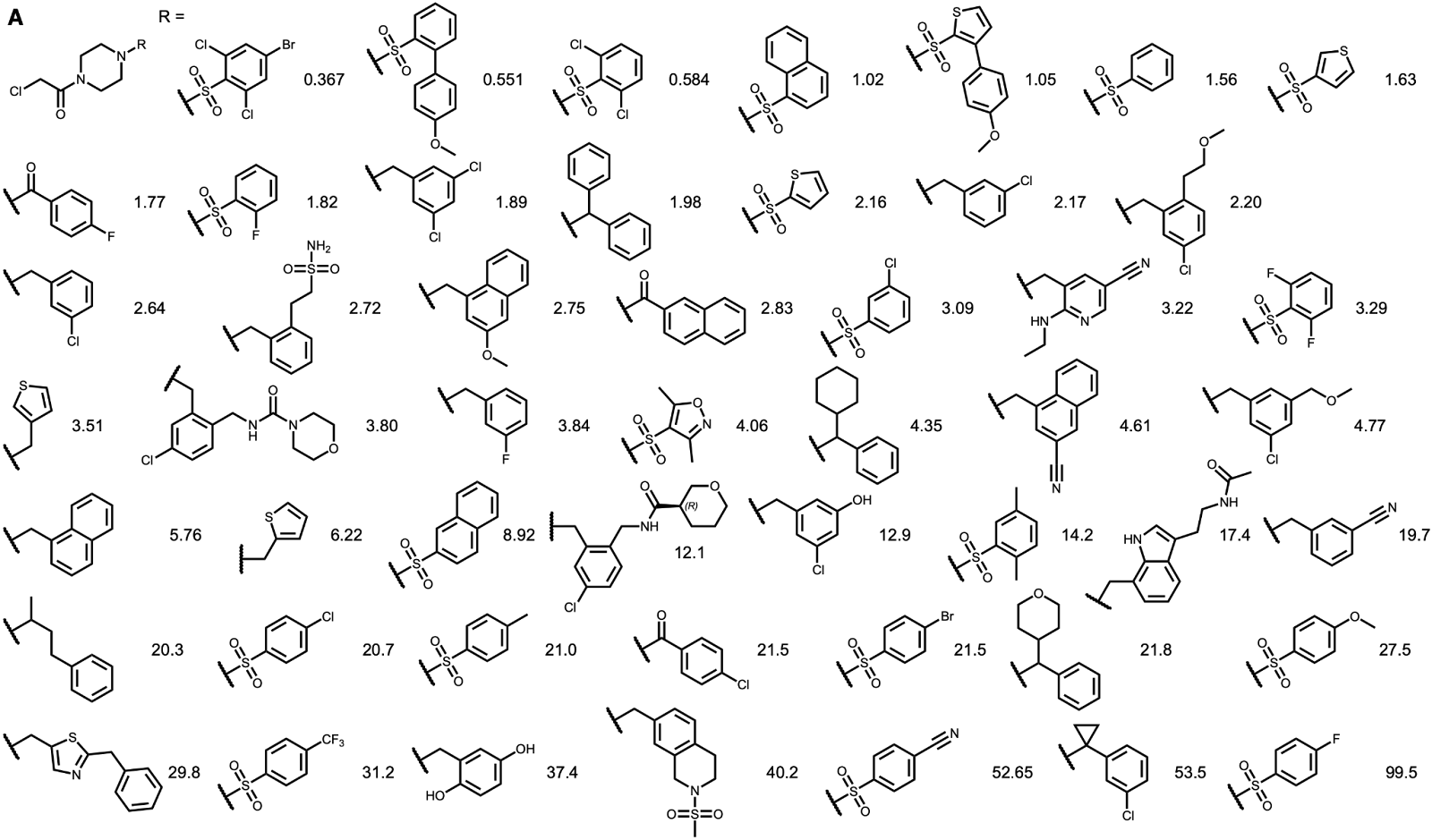

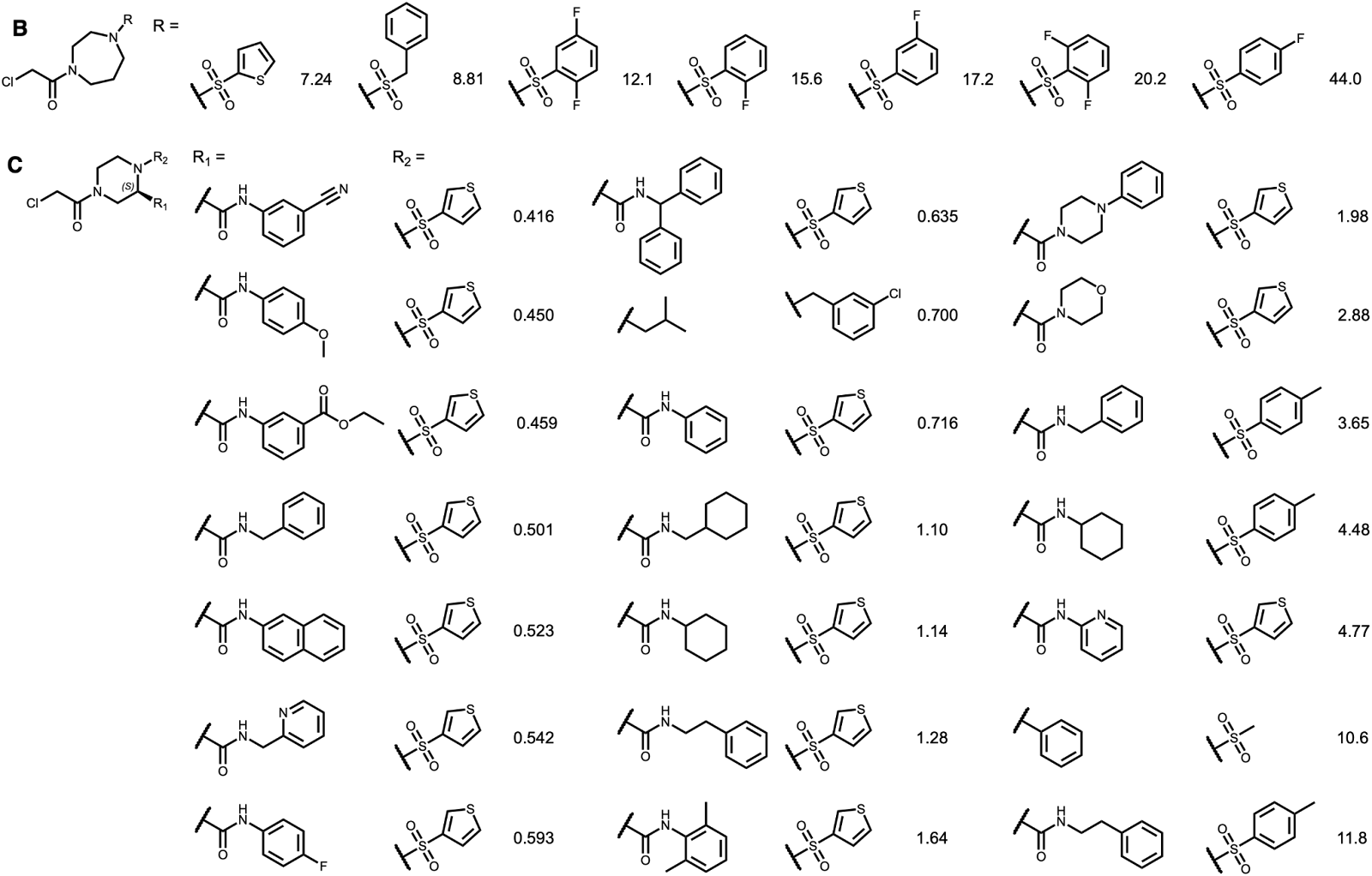

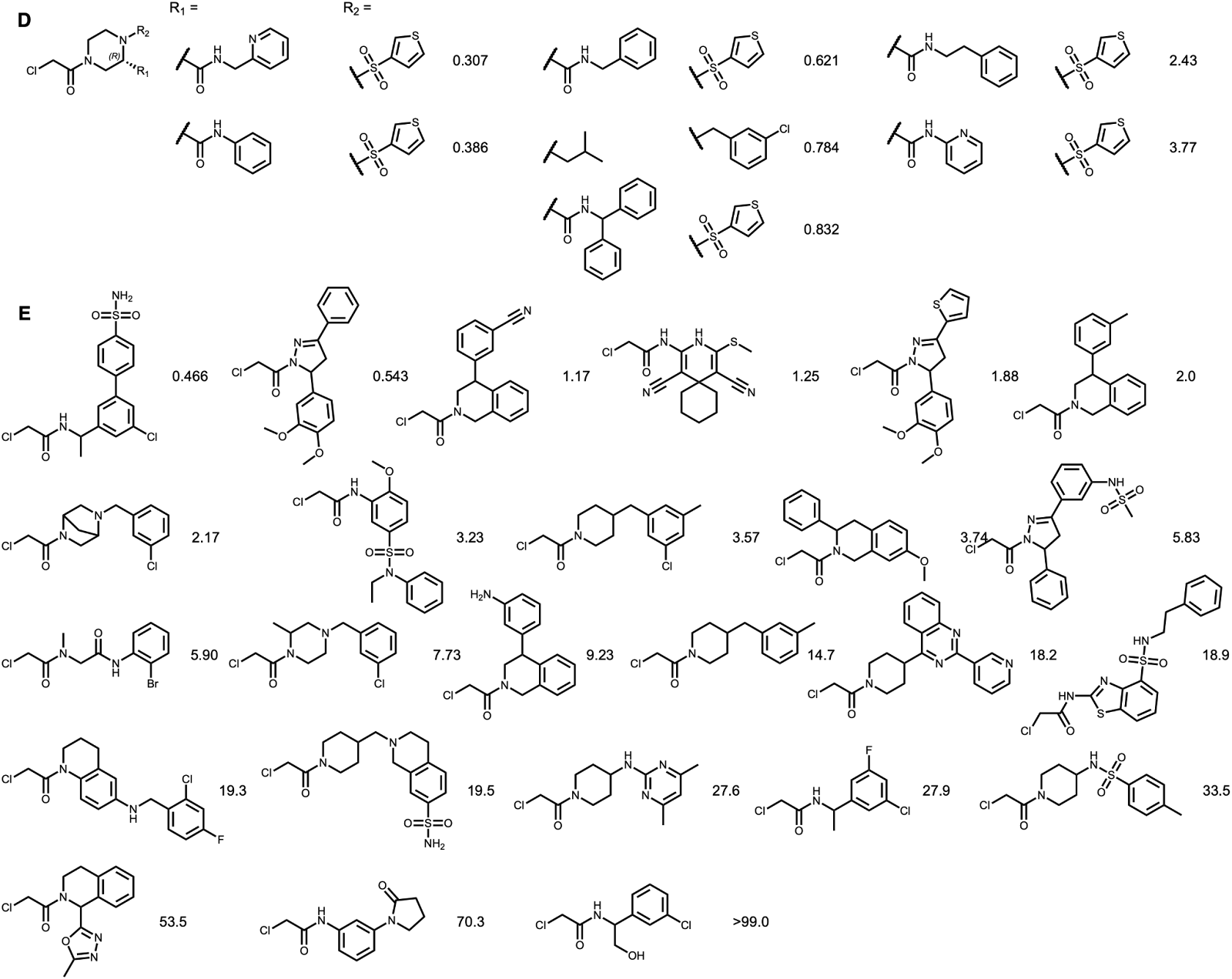

**Fig S10. Chloroacetamide fragment optimization.** Measured IC_50_ are reported in μM. **A.** Optimization of a piperazine chloroacetamide series **B.** Optimization of a diazepane  chloroacetamide series **C.** Further derivatization of the (s) isomer of the piperazine chloroacetamides **D.** Further derivatization of the (r) isomer of the piperazine chloroacetamides **E.** Singelton chloroacetamides tested.

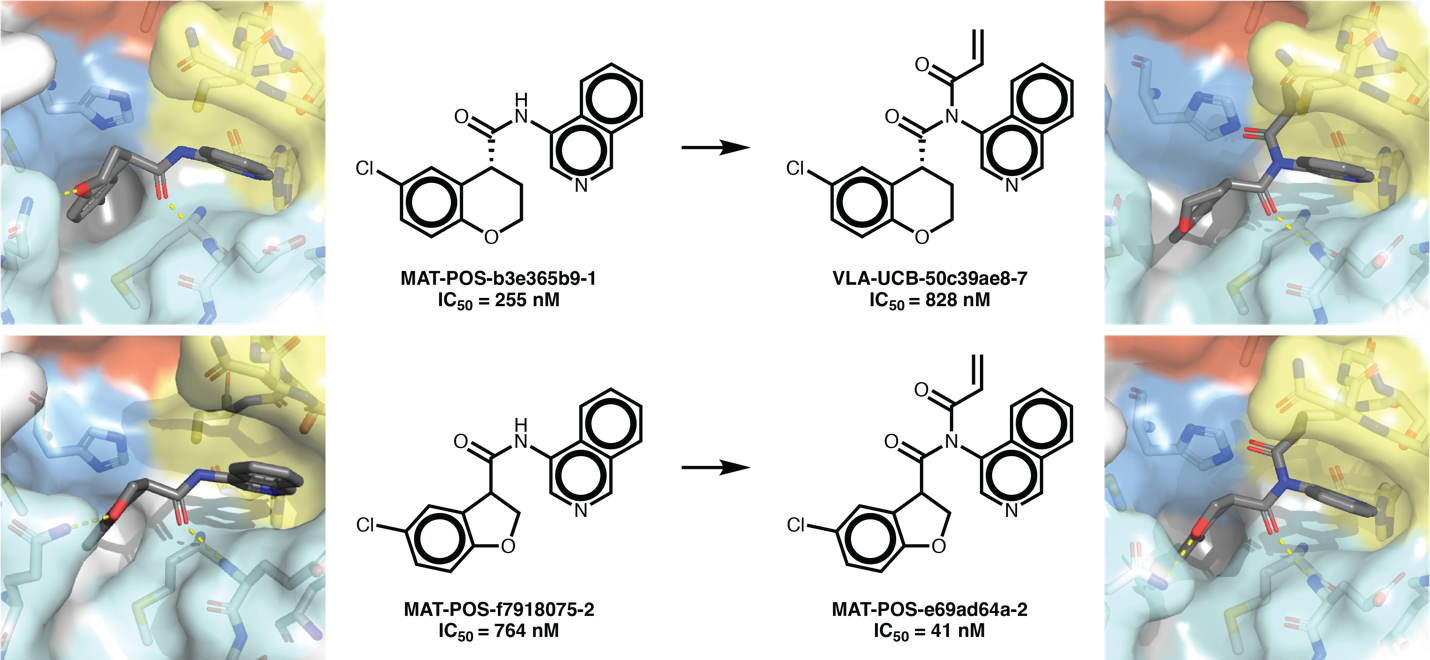

**Supplementary Figure 11: Covalentization of potent binders leads to a 40nM Mpro inhibitor.** The structure of **MAT-POS-b3e365b9-1** (top left; PDB: 7GFB) suggested that installation of an acrylamide moiety off the central amide might be able to engage covalently with the catalytic Cys145. While the acrylamide version **VLA-UCB-50c39ae8-7** proved 3-fold less potent, the co-crystal structure showed it was able indeed to form the designed covalent bond (top right; PDB: 7GJ7). Moreover, similar covalentization of a close analog (**MAT-POS-f7918075-2**; bottom left; PDB: 7GAV**)** led to **MAT-POS-e69ad64a-2** (bottom right; PDB: 7GJE) with almost 19-fold improvement in IC_50_ down to 41nM. This suggests that very slight modification of the overall binding pose can lead to dramatic rate enhancements as they place the electrophile more accurately for nucleophilic attack by the cysteine.

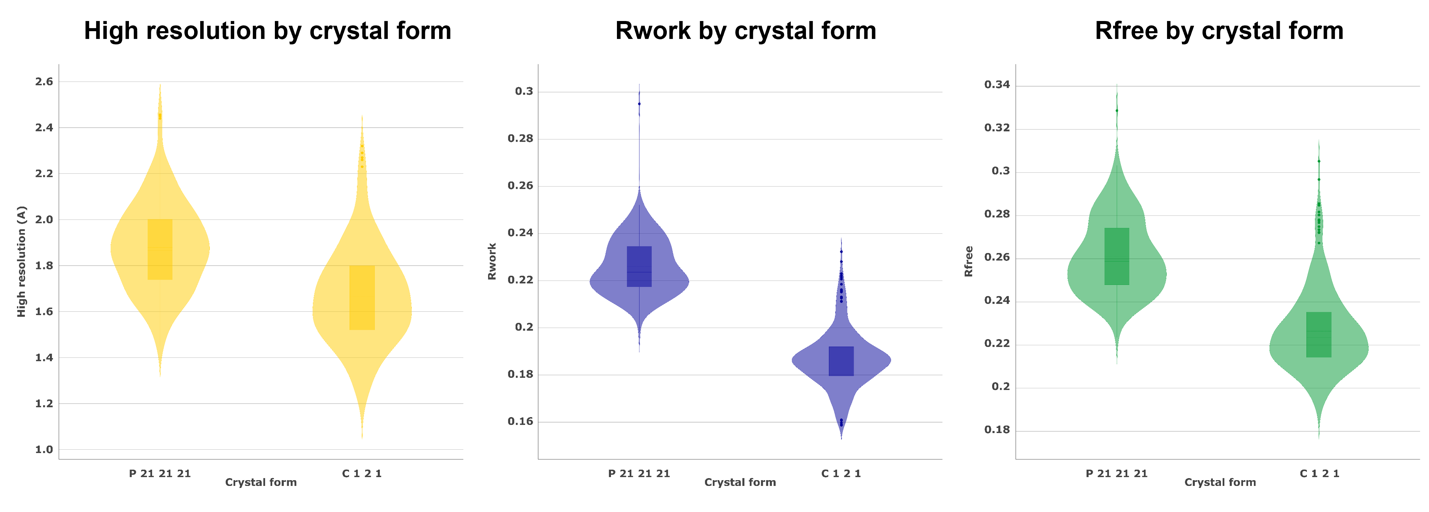

**Supplementary Figure 12: Distribution of Crystallographic statistics over collected Mpro datasets in two crystal forms.** See **Data S4** for a breakdown by structure.

**Supplementary Figure 13: Electron density maps for ligands highlighted in this manuscript.** 2FoFc maps (blue) and event maps (orange). See **Data S4** for PDB IDs.

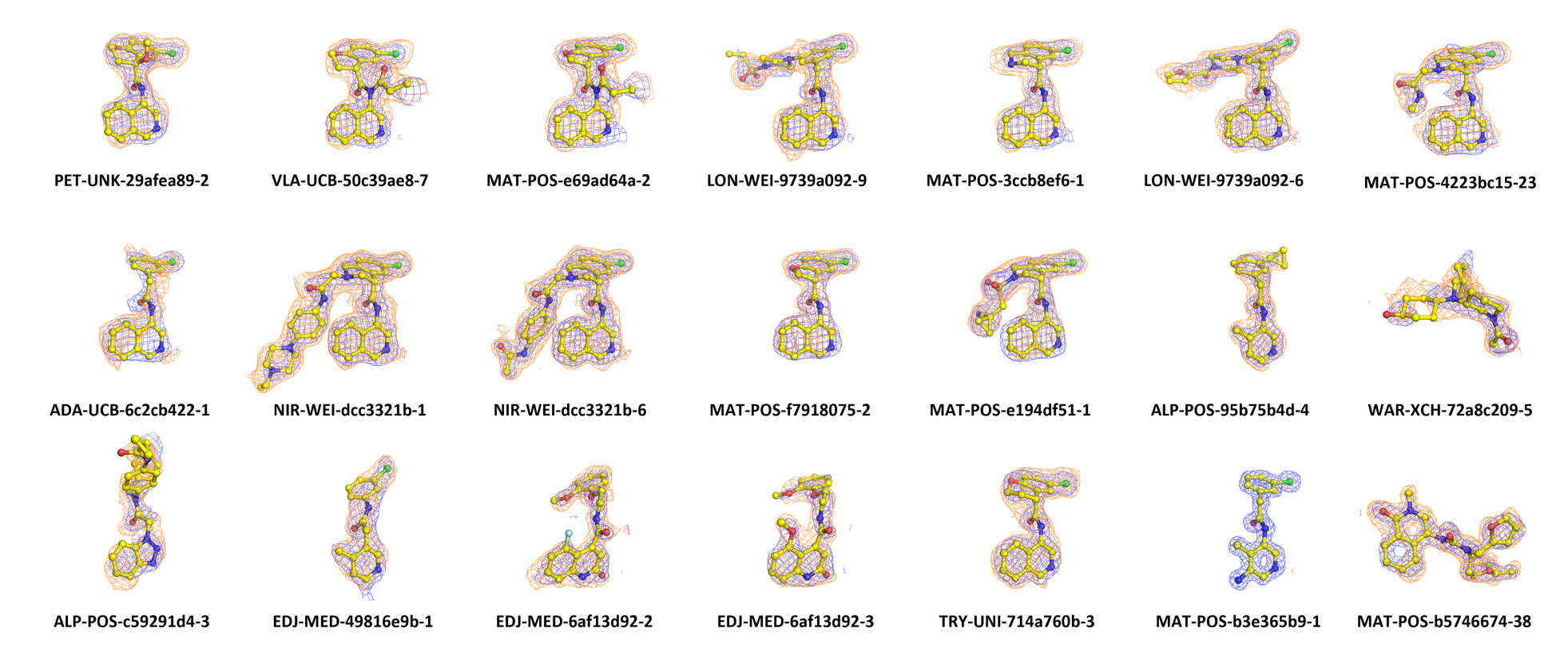

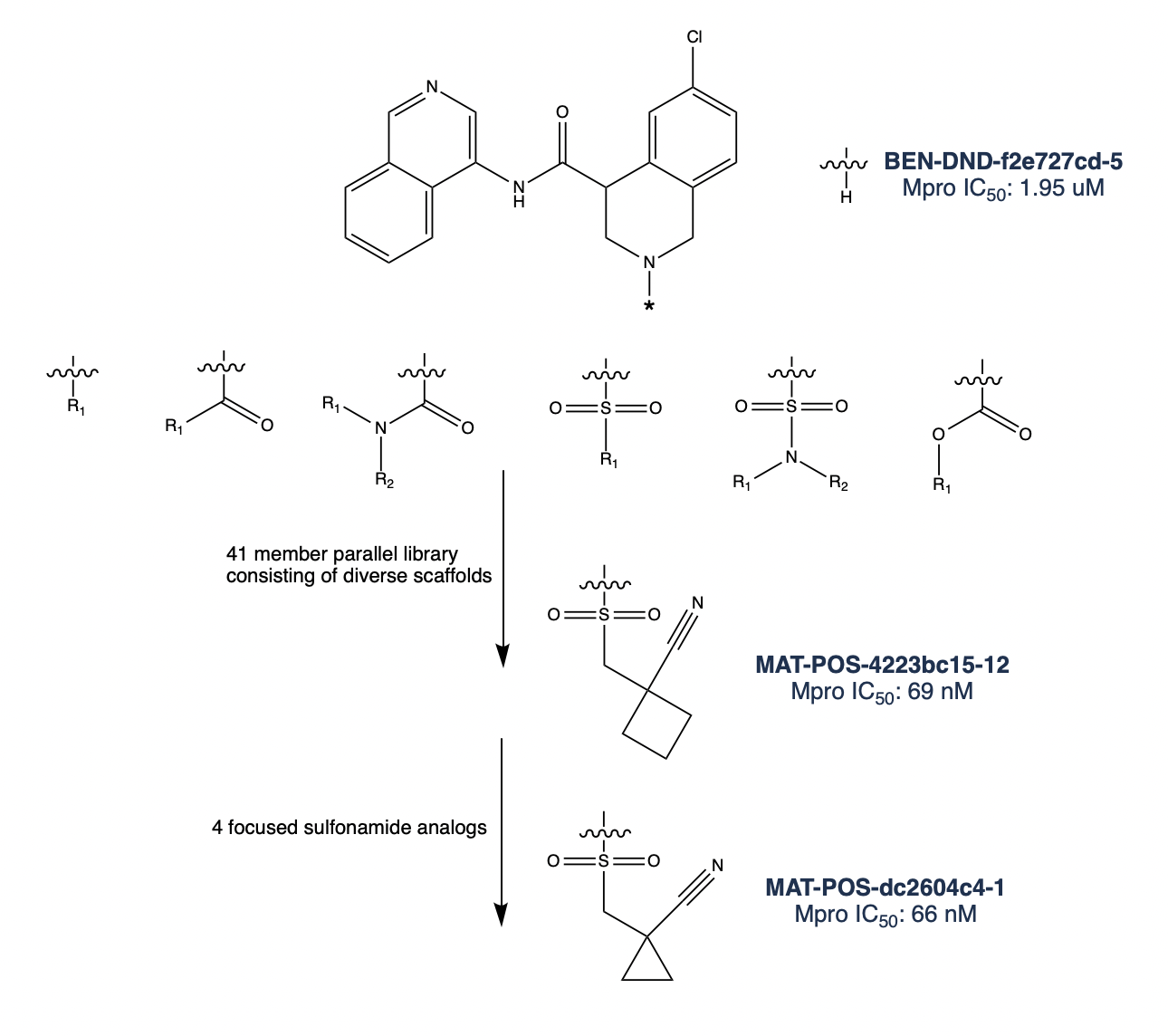

**Supplementary Figure 14:** A diverse library of potential designs accessible through robust, parallel chemistry was designed using the amine functionality of BEN-DND-f2e727cd-5 (racemate of MAT-POS-3ccb8ef6-1) as the reactive handle. The library consisted of targets accessible via standard reactions such as amide, sulfonamide, and urea couplings from large in-stock monomer classes. The large enumerated libraries were filtered down to a diverse 41 member library MAT-POS-4223bc15, which included the potent sulfonamide MAT-POS-4223bc15-12. More focused exploration of that promising compound from in-stock building blocks (MAT-POS-dc2604c4) then led to advanced compound MAT-POS-dc2604c4-1 (racemate of MAT-POS-e194df51-1).

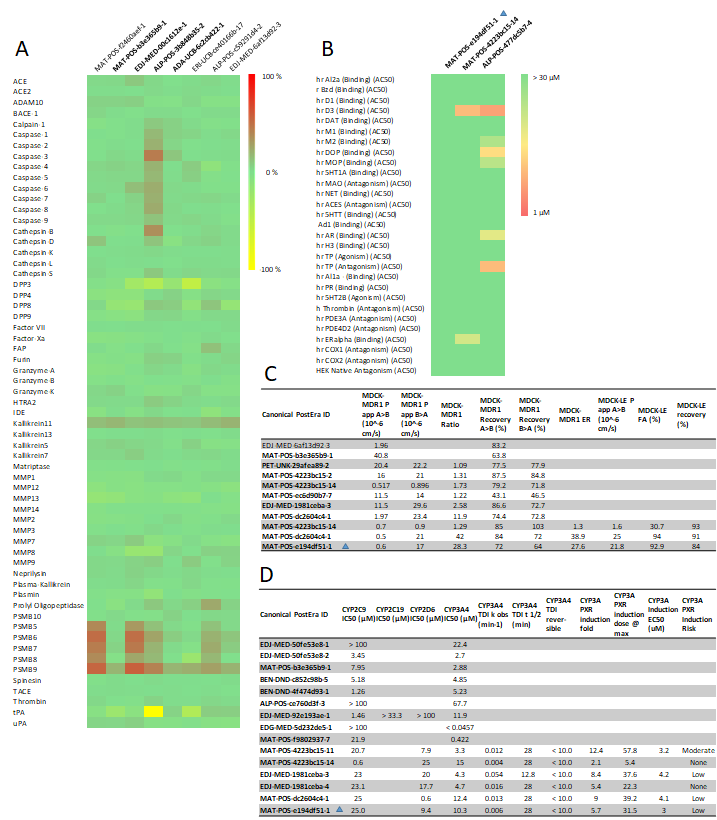

**Supplementary Figure 15: Visualization of selected safety data of the COVID Moonshot lead series.** (A) Protease enzyme selectivity measured at 100 μM (Nanosyn panel) shows high selectivity of COVID Moonshot compounds from different series against a large panel of proteases. Compounds from the aminopyridine lead series are marked in bold throughout Fig S12, with the lead compound marked by a triangle. (B) **MAT-POS-e194df51-1** shows IC50s >30 μM across the Eurofins principle panel (yellow circle), with two other compounds from the aminopyridine series also demonstrating a favourable profile. (C) Permeability data for the aminopyridine series is detailed, showing a high MDCK-MDR1 efflux ratio (ER) for **MAT-POS-e194df51-1** that suggests low brain penetrance, whilst the high MDCK-LE permeability indicates high intrinsic permeability. (D) CYP reactivity of selected COVID Moonshot compounds, showing variable engagement of Cyp3A4 across the series, but no risk of time dependent inhibition. Further, the risk of CYP3A4 induction via PXR is low.  All raw data is available as supplementary material.

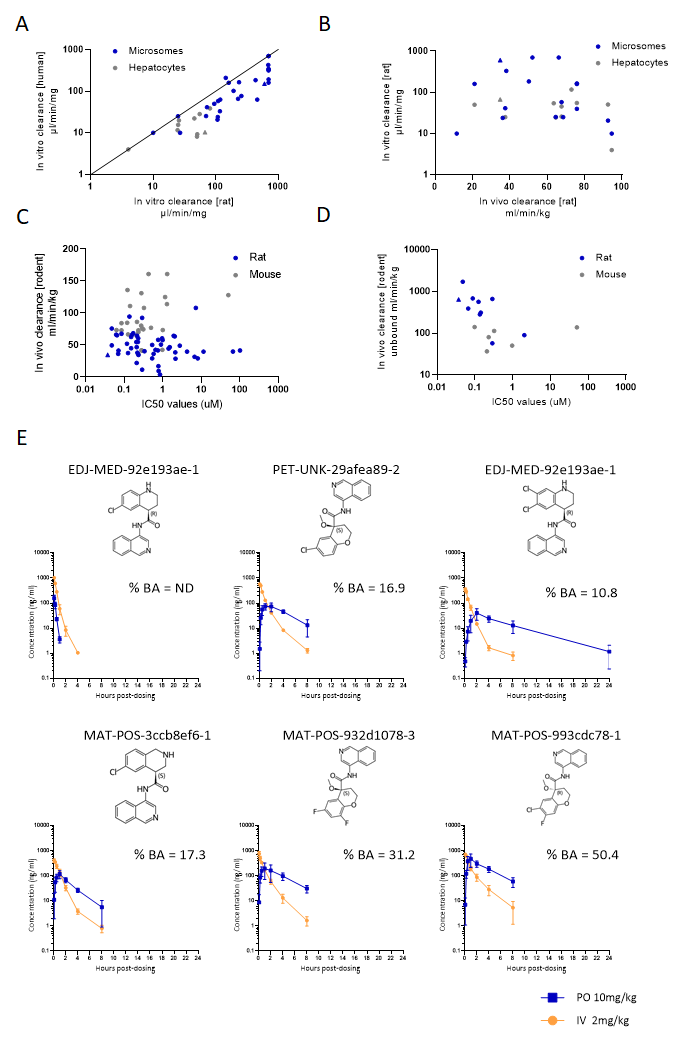

**Supplementary Figure 16: Visualisation of selected ADME, in vitro and in vivo PK data of the COVID Moonshot lead series.** (A) Both in vitro clearance measurements in microsomes (blue) and hepatocytes (grey) show a higher clearance in rats than humans. Data suggests dominant Phase 1 metabolism for the aminopyridine series. (B) Linked rat in vitro and rat in vivo clearances are depicted for microsomes and hepatocytes. (C) For all in vivo PK measurements, rat (blue) and mouse (grey) clearance is depicted in relation to fluorescence IC50 values. (D) For compounds with plasma protein binding assessed, unbound clearance is shown for rodent PK experiments. In panels A to D, **MAT-POS-e194df51-1** is marked by a triangle.  (E) Rat intravenous (IV, at 2 mg/kg) and oral (PO, at 10 mg/kg) PK experiments show the progression of oral bioavailability of the aminopyridine lead series. All raw data is available in Data S3.

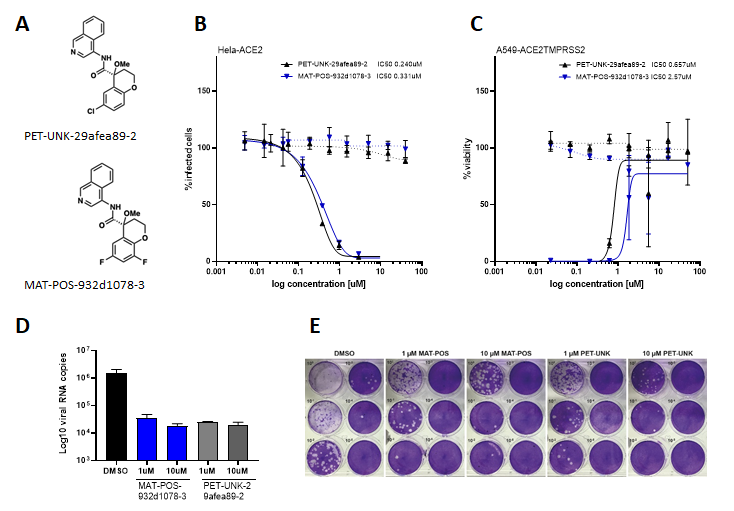

**Supplementary Figure 17: Closely related analogues of the lead compound, PET-UNK-29afea89-2 and MAT-POS-932d1078-3, demonstrate antiviral activity across different cellular antiviral assays and a kidney organoid model.  A**: shows the chemical structure of PET-UNK-29afea89-2 and MAT-POS-932d1078-3. **B:** dose-response curves of both compounds in Immunofluorescence assays in Hela-ACE2 cells, and **C**: Cytopathic Effect assays in A549-ACE2-TMPRSS2 cells. The curves also show the cytotoxicity data (dotted lines), demonstrating the lack of cytotoxic activity across all three cell lines.  **D-E**: Antiviral activity of MAT-POS-932d1078-3  and PET-UNK-29afea89-2 in kidney organoids infected with SARS-CoV-2 in the presence of 1 µM and 10 µM of compounds or DMSO as a control. **D:** Intracellular viral RNA measured by qPCRand **E**: infectious viral titers released from the apical side of the organoids at 48 hpi, measured by plaque assay on Vero E6 cells. Data in D are mean and SD of 2 biological replicates from a representative experiment of 2 independent experiments. Intracellular viral RNA levels in **D** were normalized to expression of the β-actin housekeeping gene.

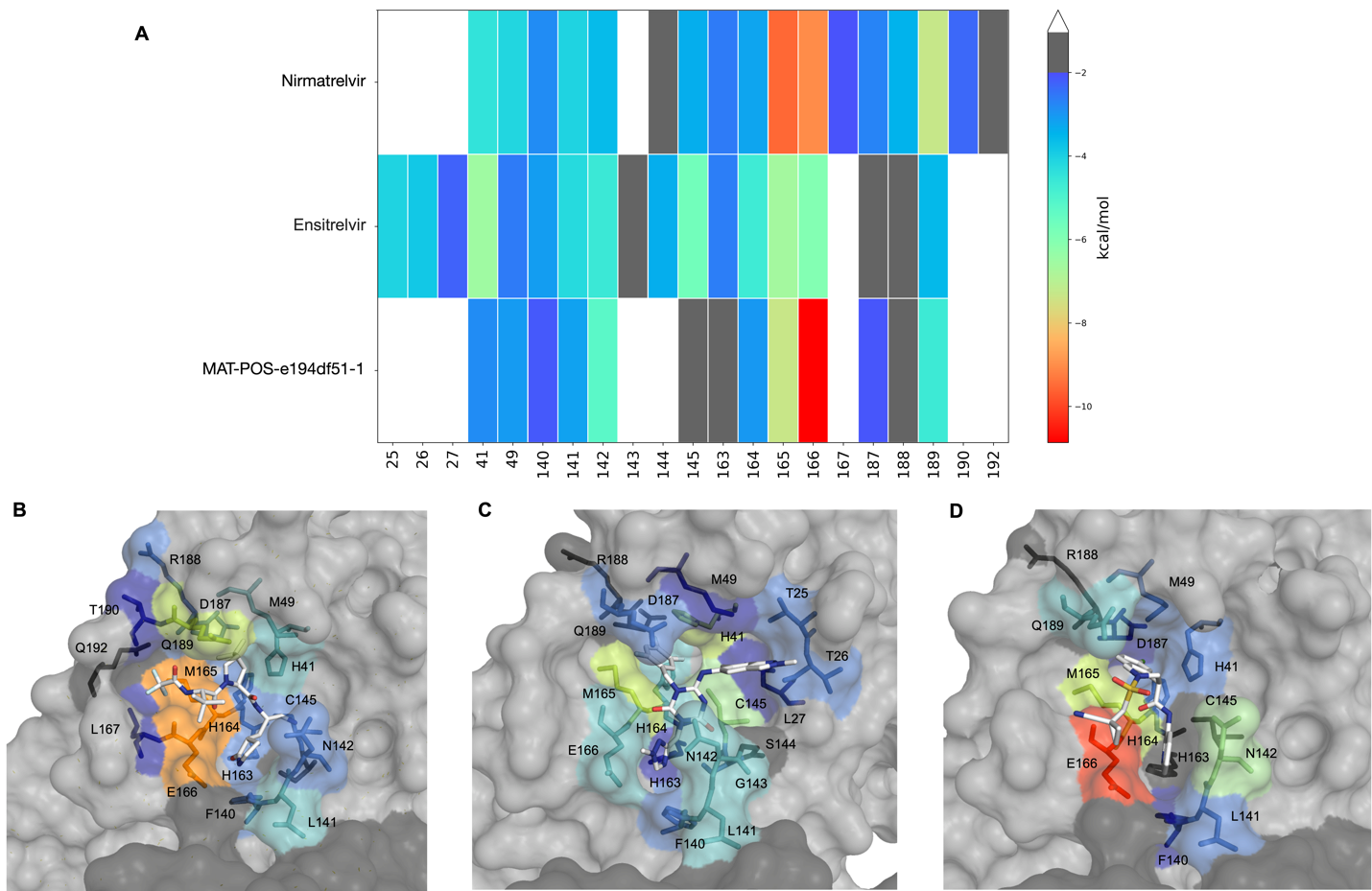

**Supplementary Figure 18**: **The interaction patterns of our lead compound MAT-POS-e194df51-1 with the Mpro binding site is distinct to known clinical antivirals nirmatrelvir and ensitrelvir (S-217622), thus likely to be active against resistant variants**. (A) The van der Waals interaction energy between the inhibitors and key residues in the Mpro binding site, and the binding poses of (B) nirmatrelvir (PDB ID: 8DZ2), (C) ensitrelvir (S-217622; PDB ID: 8DZ0) and (D) MAT-POS-e194df51-1 (PDB ID: 7GAW). The interaction energy is computed using a method previously reported in ref(*94*) and validated for hepatitis C virus protease.

**Supplementary Table 1**: **Summary crystallographic statistics for all collected datasets**

| Space group | C 1 2 1 | P 21 21 21 |
| --- | --- | --- |
| No. of datasets | 359 | 225 |
| High resolution range (A) | 1.18 - 2.84 | 1.43 - 2.44 |
| Mean high resolution (A) | 1.67 | 1.82 |
| Standard deviation high resolution (A) | 0.22 | 0.17 |
| Mean abs. dev. high resolution (A) | 0.17 | 0.13 |
| Median high resolution (A) | 1.65 | 1.83 |
| Median abs. dev. high resolution (A) | 0.14 | 0.09 |
| Mean Unique reflections (High res shell) | 92663 (1508) | 71515 (1839) |
| Mean CCHalf (High res shell) | 0.99 (0.33) | 1 (0.6) |
| Mean I/SigI (High res shell) | 8.31 (0.68) | 9.3 (1.54) |
| Mean overall completeness (High res shell) | 98.03 (93.56) | 93.5 (66.76) |
| Mean Rmerge (High res shell) | 0.12 (1.32) | 0.15 (1.25) |
| Mean multiplicity (High res shell) | 3.22 (2.76) | 6.63 (6.05) |
| Rwork range | 0.16 - 0.23 | 0.2 - 0.3 |
| Mean Rwork | 0.19 | 0.23 |
| Standard deviation Rwork | 0.01 | 0.01 |
| Mean abs. dev. Rwork | 0.01 | 0.01 |
| Median Rwork | 0.19 | 0.22 |
| Median abs. dev. Rwork | 0.01 | 0.01 |
| Rfree range | 0.19 - 0.31 | 0.22 - 0.33 |
| Mean Rfree | 0.23 | 0.26 |
| Standard deviation Rfree | 0.02 | 0.02 |
| Mean abs. dev. Rfree | 0.01 | 0.01 |
| Median Rfree | 0.22 | 0.26 |
| Median abs. dev. Rfree | 0.01 | 0.01 |

Data S1. (separate file)

Summary of all collected biochemical data.

Data S2. (separate file)

Summary of designs uploaded to the COVID Moonshot platform.

Data S3. (separate file)

Summary of nanomole high throughput optimization.

Data S4. (separate file)

Summary of crystallographic and refinement statistics

Data S5. (separate file)

Summary of ADMET and PK data.

Data S6. (separate file)

Summary of all cellular anti-viral data produced in this project.

Data S7. (separate file)

The Covid Moonshot consortium member list
